# Laboratory evolution of a *Saccharomyces cerevisiae* x *S. eubayanus* hybrid under simulated lager-brewing conditions: genetic diversity and phenotypic convergence

**DOI:** 10.1101/476929

**Authors:** Arthur R. Gorter de Vries, Maaike A. Voskamp, Aafke C. A. van Aalst, Line H. Kristensen, Liset Jansen, Marcel van den Broek, Alex N. Salazar, Nick Brouwers, Thomas Abeel, Jack T. Pronk, Jean-Marc G. Daran

## Abstract

*Saccharomyces pastorianus* lager-brewing yeasts are domesticated hybrids of *S. cerevisiae* x *S. eubayanus* that display extensive inter-strain chromosome copy number variation and chromosomal recombinations. It is unclear to what extent such genome rearrangements are intrinsic to the domestication of hybrid brewing yeasts and whether they contribute to their industrial performance. Here, an allodiploid laboratory hybrid of *S. cerevisiae* and *S. eubayanus* was evolved for up to 418 generations on wort under simulated lager-brewing conditions in six independent sequential batch bioreactors. Characterization of 55 single-cell isolates from the evolved cultures showed large phenotypic diversity and whole-genome sequencing revealed a large array of mutations. Frequent loss of heterozygosity involved diverse, strain-specific chromosomal translocations, which differed from those observed in domesticated, aneuploid *S. pastorianus* brewing strains. In contrast to the extensive aneuploidy of domesticated *S. pastorianus* strains, the evolved isolates only showed limited (segmental) aneuploidy. Specific mutations could be linked to calcium-dependent flocculation, loss of maltotriose utilisation and loss of mitochondrial activity, three industrially relevant traits that also occur in domesticated *S. pastorianus* strains. This study indicates that fast acquisition of extensive aneuploidy is not required for genetic adaptation of *S. cerevisiae* x *S. eubayanus* hybrids to brewing environments. In addition, this work demonstrates that, consistent with the diversity of brewing strains for maltotriose utilization, domestication under brewing conditions can result in loss of this industrially relevant trait. These observations have important implications for the design of strategies to improve industrial performance of novel laboratory-made hybrids.

## Introduction

*Saccharomyces* yeasts are popular eukaryotic models for studying genome hybridisation, chromosome (mis)segregation and aneuploidy (Botstein, et al. 1997; Sheltzer, et al. 2011). The genus *Saccharomyces* arose between 10 and 20 million years ago and currently comprises eight described species, as well as interspecies hybrids (Liti, et al. 2006; Hittinger 2013; Naseeb, et al. 2017). Absence of a prezygotic barrier between *Saccharomyces* species facilitates hybridization, although spore viabilities of the resulting hybrids is typically well below 10 % (Liti, et al. 2006; Hittinger 2013; Naseeb, et al. 2017). Several interspecies *Saccharomyces* hybrids are tightly associated with domestication in industrial processes. *S. pastorianus* lager-brewing yeasts are domesticated *S. cerevisiae* x *S. eubayanus* hybrids (Libkind, et al. 2011). Double and triple hybrids between *S. cerevisiae*, *S. kudriavzevii* and *S. uvarum* are closely associated with wine fermentation (González, et al. 2006; Querol and Bond 2009; Marsit and Dequin 2015). *S. bayanus* cider fermentation yeasts are domesticated *S. uvarum* X *S. eubayanus* hybrids (Naumov, et al. 2001). Reconstruction of the corresponding *Saccharomyces* hybrids in the laboratory showed improved performance, relative to the parental species. For example, laboratory-made *S. cerevisiae* x *S. eubayanus* hybrids combined sugar utilisation characteristics of *S. cerevisiae* and the superior performance at low temperatures of *S. eubayanus* (Hebly, et al. 2015; Krogerus, et al. 2017). Similarly, hybrids of *S. cerevisiae*, *S. kudriavzevii* and *S. uvarum* combined traits of their parental species relevant to industrial wine fermentation, such as flocculence, sugar utilisation kinetics, stress tolerance and aroma production (Coloretti, et al. 2006; Lopandic, et al. 2016).

The relevance of laboratory hybridization of *Saccharomyces* species extends beyond reconstruction of existing, domesticated hybrids. The ability of hybridisation to generate extensive phenotypic diversity has raised interest in the development of novel *Saccharomyces* hybrids for specific industrial processes (Krogerus, et al. 2017). For example, an *S. cerevisiae* × *S. paradoxus* hybrid produced high concentrations of aromatic compounds that are of interest for wine making (Bellon, et al. 2011). Hybrids between *S. cerevisiae* and *S. arboricola* or *S. mikatae* were able to utilize the sugars in wort at low temperatures and produced particularly aromatic beer (Nikulin, et al. 2017). Laboratory hybrids of *S. cerevisiae* and *S. kudriavzevii* or *S. mikatae* yielded xylose-consuming strains with high inhibitor tolerance for 2^nd^ generation biofuel production (Peris, et al. 2017).

The alloeuploid genomes of laboratory hybrids of *Saccharomyces* species strongly differ from the extremely aneuploidy genomes of the domesticated strains used in traditional industrial processes. For example, the genomes of *S. pastorianus* lager-brewing yeasts contain between 45 and 79 chromosomes (Van den Broek, et al. 2015; Okuno, et al. 2016), a degree of aneuploidy that is not observed elsewhere in the *Saccharomyces* genus (Gorter de Vries, Pronk, et al. 2017). The current consensus is that all *S. pastorianus* strains have evolved from a single, common ancestor *S. cerevisiae* x *S. eubayanus* hybrid by centuries of domestication and strain improvement in brewing-related environments (Okuno, et al. 2016). However, it remains unclear when and how domestication resulted in the extensive chromosome copy number variations and phenotypic diversity of current *S. pastorianus* strains.

Hybrid genomes have a well-documented increased tendency to become aneuploid due to an increased rate of chromosome missegregation during mitosis and/or meiosis (Chambers, et al. 1996; Liti, et al. 2006). Aneuploidy reduces the efficiency of sporulation and can thereby complicate genetic modification, impeding breeding and targeted strain improvement (Santaguida and Amon 2015; Gorter de Vries, de Groot, et al. 2017). In evolutionary contexts, aneuploidy is generally seen as a transient adaptation mechanism, whose positive impacts are eventually taken over by more parsimonious mutations (Yona, et al. 2012). When grown mitotically, sporulated hybrid strains were prone to further chromosome missegregation resulting in more extensive chromosome copy number variations (Lopandic, et al. 2016). Even genomes of *Saccharomyces* hybrids that had not undergone meiosis displayed increased rates of mitotic chromosome missegregation during mitosis (Delneri, et al. 2003). Indeed, when evolved in lignocellulosic hydrolysates, cultures of *S. cerevisiae X S. kudriavzevii* and *S. cerevisiae* x *S. mikatae* hybrids exhibited segmental and full-chromosome aneuploidy after only 50 generations (Peris, et al. 2017). Similarly, when evolved under wine fermentation conditions, *S. cerevisiae* x *S. kudriavzevii* hybrids displayed extensive genome reorganisations that led to a significant reduction of their genome content (Pérez Través, et al. 2014).

Genetic instability of hybrid genomes could be detrimental to stable, robust industrial performance. Therefore, to assess industrial applicability of new hybrids generated in the laboratory, it is important to determine their genome stability under industrially relevant conditions. Moreover, laboratory evolution under simulated industrial conditions can increase understanding of the selective pressures that shaped the genomes of domesticated microorganisms (Bachmann, et al. 2012; Gibbons, et al. 2012; Gibbons and Rinker 2015).

The goal of the present study was to investigate how a previously constructed allodiploid *S. cerevisiae* x *S. eubayanus* hybrid (Hebly, et al. 2015) evolves under simulated lager-brewing conditions, with a specific focus on genome dynamics and on acquisition or loss of brewing-related phenotypes. To mimic successive lager beer fermentation processes, the hybrid strain was subjected to sequential batch cultivation on industrial wort, in six independent bioreactor setups. After up to 418 generations, the genotypic and phenotype diversity generated in these laboratory evolution experiments was analysed by characterization of 55 single-cell isolates. After whole-genome resequencing of each isolate using 150 bp paired-end reads, sequence data were mapped to high-quality reference genomes of the parental strains to identify genomic changes. Phenotypic analysis of the isolates focused on the ability to utilise maltotriose, flocculation and the respiratory capacity. We interpreted these results in the context of the domestication history of *S. pastorianus* brewing strains as well as in relation to genome stability and industrial application of newly generated *Saccharomyces* hybrids.

## Results

### Simulating domestication under lager-brewing conditions in sequential batch bioreactors

Industrial lager brewing involves batch cultivation of *S. pastorianus* on wort, an extract from malted barley, at temperatures between 7 and 15 °C. After a brief initial aeration phase to enable oxygen-dependent biosynthesis of unsaturated fatty acids and sterols (Andreasen and Stier 1953; Gibson, et al. 2007), brewing fermentations are not aerated or stirred, leading to anaerobic conditions during the main fermentation (Briggs, et al. 2004). To simulate domestication under industrial lager-brewing conditions, a laboratory evolution regime was designed in which the laboratory-made *S. cerevisiae* x *S. eubayanus* hybrid IMS0408 was grown at 12 °C in sequential batch bioreactors on industrial wort. As in industrial brewing, each cultivation cycle was preceded by an aeration phase, after which cultures were incubated without sparging or stirring until a decline of the CO_2_ production indicated a cessation of sugar consumption. The bioreactors were then partially emptied, leaving 7 % of the culture volume as inoculum for the next aeration and fermentation cycle, which was initiated by refilling the reactor with sterile wort (Figure 1A and 1B). To mimic the low sugar concentrations during early domestication of *S. pastorianus* (Meussdoerffer 2009), parallel duplicate experiments at 12 °C were not only performed with full-strength 17 °Plato wort (‘<u>H</u>igh <u>G</u>ravity’, experiments HG12.1 and HG12.2), but also with three-fold diluted wort (‘<u>L</u>ow <u>G</u>ravity’, experiments LG12.1 and LG12.2). To enable a larger number of generations during 4 months of operation, additional duplicate experiments on three-fold diluted wort were performed at 30 °C (LG30.1 and LG30.2).

**Figure 1:**
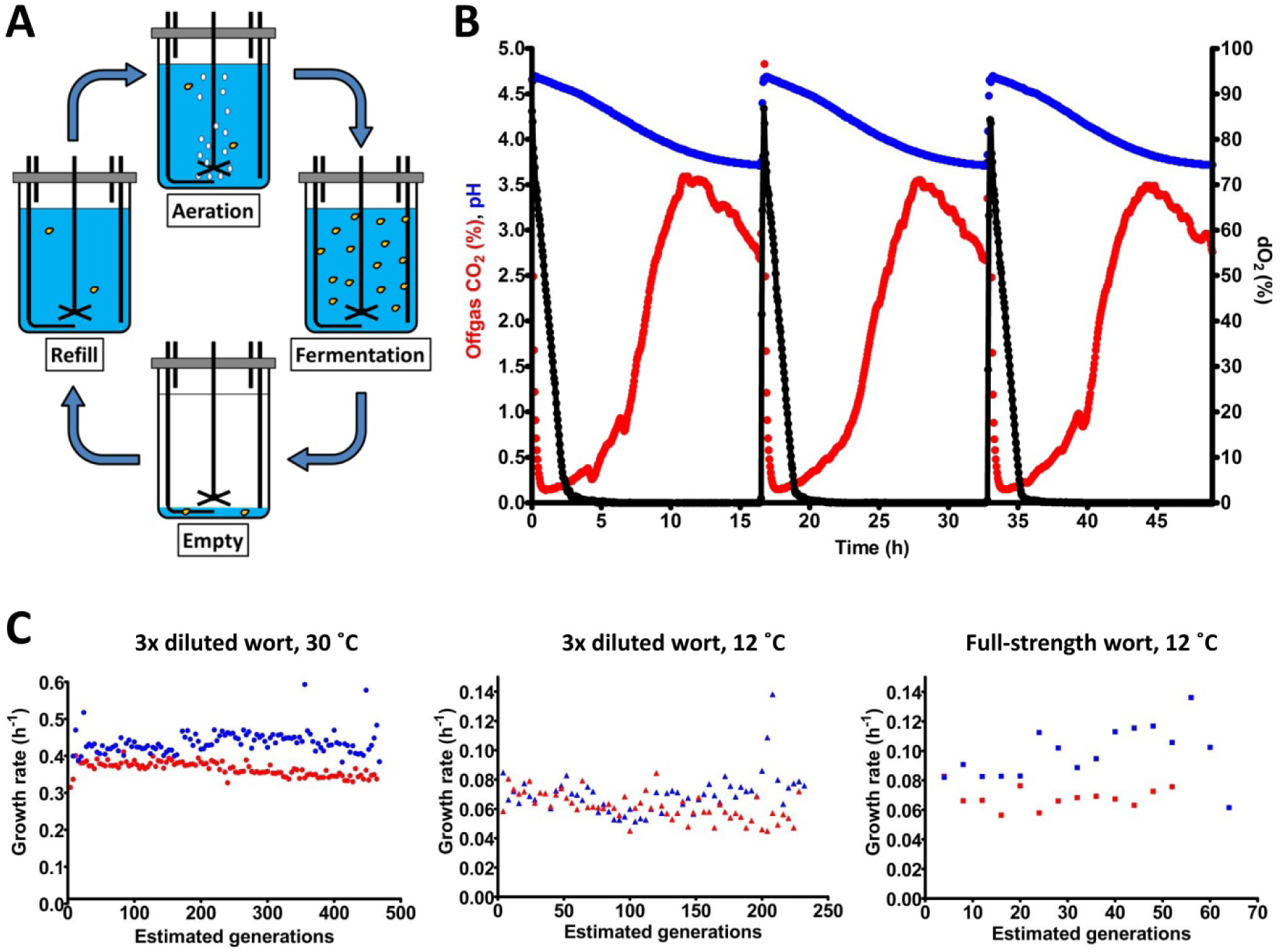
Laboratory evolution mimicking the domestication of lager-brewing yeast. The *S. cerevisiae* x *S. eubayanus* laboratory hybrid IMS0408 was grown in duplicate sequential batch bioreactors in 3-fold diluted wort at 30 °C (LG30.1 and LG30.2) and at 12 °C (LG12.1 and LG12.2), and in full-strength wort at 12 °C (HG12.1 and HG12.2). (A) Experimental design for simulated sequential lager-beer brewing cycles. Each cycle consisted of four phases: (i) (re)filling of the fermenter with fresh medium up to a total volume of 100 mL, (ii) aeration at 200 mL/min while stirring at 500 RPM, (iii) a batch fermentation phase without sparging or stirring, while flushing the bioreactor headspace with N_2_ to enable accurate analysis of CO_2_ production and (iv) removal of broth, leaving 7 mL to inoculate the next cycle. (B) Fermentation profiles of three consecutive cycles from experiment LG30.1, performed at 30 °C in 3-fold diluted wort. Percentage of CO_2_ in the off gas, culture pH and dissolved oxygen (dO_2_) concentration are indicated by red, blue and black symbols, respectively. Due to the lack of stirring and sparging, CO_2_ was slowly released by the medium; emptying of the reactor was initiated when the offgas CO_2_ concentration dropped to 70 % of its initial value as off-line analyses indicated that, at this point, all fermentable sugars had been consumed (C) Specific growth estimated from CO_2_ production profiles during each cycle of the evolution lines. LG30.1 (blue circles) and LG30.2 (red circles) were grown on 3-fold diluted wort at 30 °C; LG12.1 (blue triangles) and LG12.2 (red triangles) were grown on 3-fold diluted wort at 12 °C. HG12.1 (blue squares) and HG12.2 (red squares) were evolved in full-strength wort at 12 °C. Since lack of sparging and stirring precluded exact estimates of specific growth rates, the calculated values should be taken as indicative.

Concentration of the wort and temperature strongly affected the length of the fermentation cycles, which was 17 h for LG30.1 and LG30.2, 93 h for LG12.1 and LG12.2 and 205 h for HG12.1 and HG12.2. Experiments LG30.1 and LG30.2 involved 117 and 118 batch cycles, respectively, LG12.1 and LG12.2 covered 58 and 57 cycles, respectively, and HG12.1 and HG12.2 covered 13 and 16 cycles, respectively. At the inoculum size of 7 % of the total culture volume, each cycle corresponded to approximately 4 generations. Specific growth rates, estimated from CO_2_ production rates during the exponential growth phase of the batch cycles, were not significantly different during the first and the last five cycles of each experiment (Student’s t-test, p > 0.05). Average specific growth rates were 0.35 ± 0.02 h^−1^ for LG30.1, 0.42 ± 0.03 h^−1^ for LG30.2, 0.070 ± 0.013 h^−1^ for LG12.1, 0.062 ± 0.009 h^−1^ for LG12.2, 0.068 ± 0.007 h^−1^ for HG12.1 and 0.098 ± 0.018 h^−1^ for HG12.2. While the initial specific growth rate was clearly higher at 30 °C than at 12 °C, initial growth rates on diluted and full-strength wort were not significantly different. However, CO_2_ production from sugars continued much longer in full-strength wort. During brewing fermentation, depletion of nitrogen sources and oxygen limit biomass formation. Complete sugar conversion therefore depends on growth-independent alcoholic fermentation which, apparently, was much slower in cultures grown on full-strength wort. At the end of each evolution experiment, culture samples were streaked on YPD agar and 5 single colonies were isolated for each culture. For experiments LG12.1 and LG12.2, isolates were also made from intermediate samples after 29 cycles. Evolution lines LG30.2 and LG12.2 developed flocculence and isolates from these lines had two distinct colony morphologies: about half of the colonies were elevated and conically-shaped, while the other colonies shared the flat morphology of IMS0408. For each of these lines, five random colonies of each morphology were selected.

### Prolonged growth under simulated brewing conditions did not cause large ploidy changes

Six independent sequential batch fermentation experiments under simulated brewing conditions, covering 52 to 468 generations, yielded 55 isolates. Staining with the DNA-binding fluorescent dye SYTOX Green and flow cytometry indicated genome sizes of the isolates between 17.6 and 23.5 Mbp (Supplementary table S1). These values did not differ significantly from the 21.3 ± 1.9 Mbp genome size measured for the parental laboratory hybrid IMS0408 and therefore indicated the absence of large changes in genome content such as whole-genome duplications. For a detailed genotypic analysis, the genomes of the 55 isolates were sequenced using 150 bp pair-end reads with 101- to 189-fold coverage. A high quality-reference genome was constructed by combining the chromosome-level contigs from assemblies of CEN.PK113-7D and CBS12357 generated with nanopore technology, including mitochondrial genome sequences (Salazar, et al. 2017; Brickwedde, et al. 2018).

Copy number analysis revealed whole-chromosome aneuploidies in only 5/55 isolates (Table 1). Relative to strain IMS0408, the total chromosome number of the isolates had not changed by more than one. Isolate IMS0556 (LG30.1) had gained a copy of *Sc*CHRVIII, IMS0560 (LG30.1) had gained a copy of *Se*CHRX, IMS0565 (LG30.2) had lost *Sc*CHRXIV and gained a copy of *Se*CHRXIV, IMS0595 (LG12.1) had gained a copy of *Se*CHRVIII and IMS0606 (LG12.2) had lost a copy of *Se*CHRVIII. Read alignments to mitochondrial genome sequences were absent from 14/55 isolates, while 1 isolate showed only a partial alignment, indicating complete (ρ^−^) or partial loss (ρ^0^) of the mitochondrial genome in 15/55 strains. Loss of respiratory competence was confirmed by the observation that these 15 isolates, in contrast to IMS0408 and isolates containing a full mitochondrial genome, were unable to grow on YP-ethanol (Supplementary Figure S1).

**Table 1:**
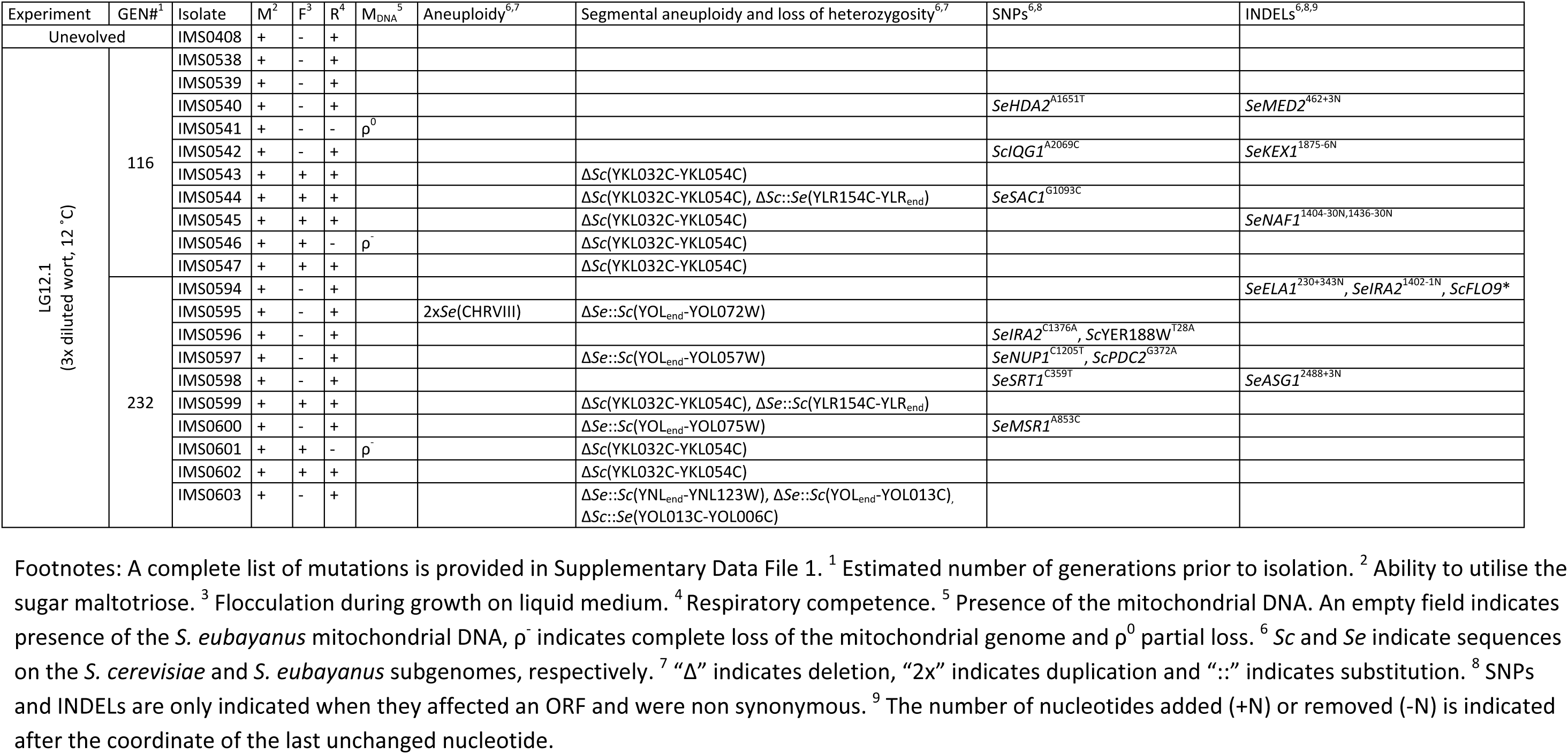
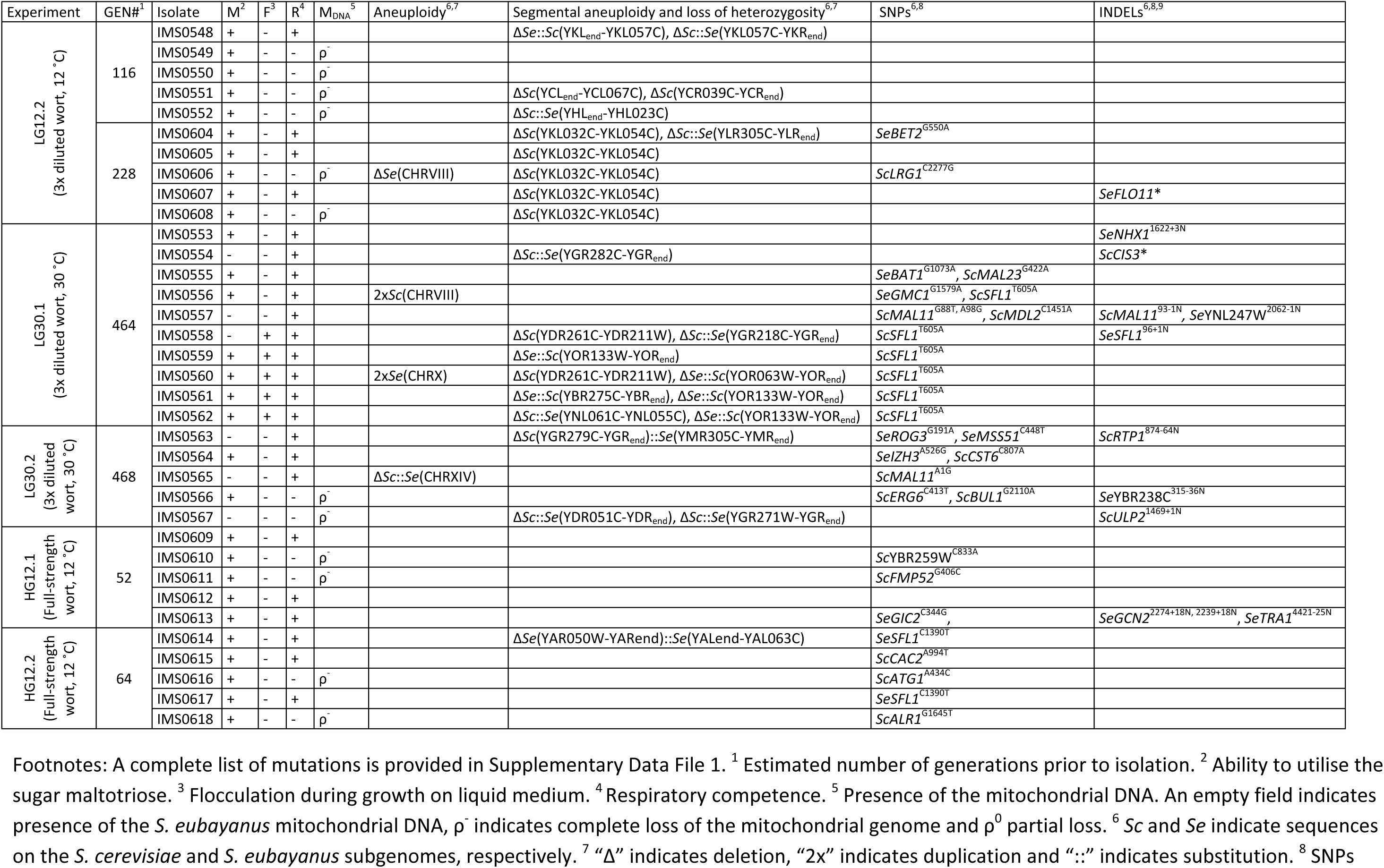
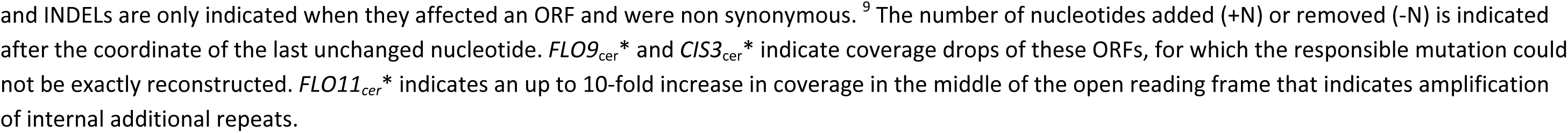
Overview of phenotypic and genotypic changes in isolates obtained after laboratory evolution of the allodiploid laboratory hybrid IMS0408 under simulated lager-brewing conditions.

### Chromosomal recombinations frequently caused loss of heterozygosity

Of the 55 evolved isolates, 29 displayed segmental copy number changes. In total, 20 of the 32 chromosomes in strain IMS0408 were affected in at least one isolate. Of the 55 evolved isolates, 24 showed chromosome segments with increased copy number and 41 showed chromosome segments with decreased copy number (Table 1, Figure 2). 17 internal recombinations resulting in deletions were observed: Δ*Sc*(YKL032C-YKL054C) occurred in 13 strains, Δ*Sc*(YDR261C-YDR211W) occurred in two strains, and Δ*Sc*(YCL_end_-YCL067C) and Δ*Sc*(YCR039C-YCR_end_) occurred together in one strain. The internal recombinations Δ*Sc*(YKL032C-YKL054C) and Δ*Sc*(YDR261C-YDR211W) both resulted in loss of the sequence between the recombination sites. The recombination occurred between *IXR1* and *DEF1* in Δ*Sc*(YKL032C-YKL054C) and between Ty-transposons for Δ*Sc*(YDR261C-YDR211W). Finally, the concurrent loss of *Sc*(YCL_end_-YCL067C) and *Sc*(YCR039C-YCR_end_) indicated loss of both ends of *Sc*CHRIII, including telomeres, consistent with recombination into a circular chromosome III at the HMLALPHA2 and MATALPHA2 loci (Newlon, et al. 1991).

**Figure 2:**
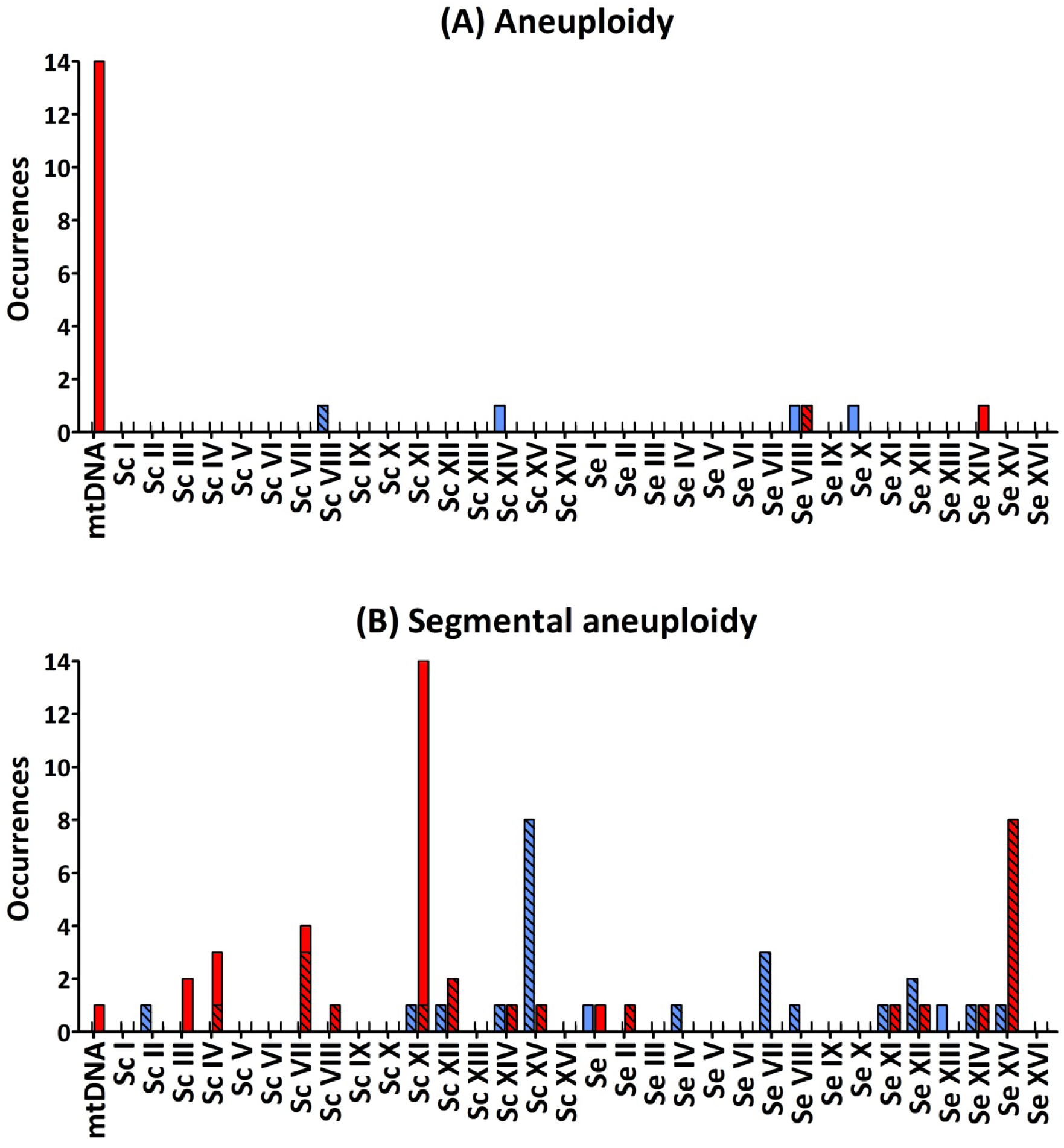
Total number of occurrences of whole-chromosome (A) and segmental (B) aneuploidy for each chromosome of IMS0408 among 55 isolates obtained after laboratory evolution under simulated lager fermentation conditions. For each chromosome, loss of genetic material is indicated in red and duplicated genetic material is indicated in blue. Loss or duplication of *S. cerevisiae* or *S. eubayanus* genetic material which was coupled with duplication or loss of the corresponding region of the other subgenome, is indicated by checked bars. *S. eubayanus* harbours two translocations relative to *S. cerevisiae:* between chromosomes II and IV, and between chromosomes VIII and XV. For simplicity, copy number affecting these regions were allocated based on the *S. cerevisiae* genome architecture.

The remaining 24 chromosome-segment duplications and losses reflected inter-chromosomal recombinations: one chromosomal region was replaced by an additional copy of another chromosomal region by a non-conservative recombination. The recombinations Δ*Sc*(YGR279C-YGR_end_)::*Se*(YMR305C-YMR_end_) and Δ*Se*(YAR050W-YAR_end_)::*Se*(YAL_end_-YAL063C) occurred between highly similar genes; the paralogs *SCW4* and *SCW10,* and *FLO1* and *FLO9*, respectively. In the remaining 22 cases, recombination occurred between homologous genes of each subgenome. No copy-number conservative chromosome translocations were identified.

Of the 26 observed recombinations, 23 occurred inside ORFs, and thus resulted in chimeric genes (Table 2). The homology between ORFs involved in recombinations varied from <70 % to 100 %, with a median homology of 82.41 %. Chimeric ORFs were reconstructed by extracting reads paired to the other chromosome from the sequencing data and using them for a local assembly. This approach allowed for identification of the recombination site at a resolution that, depending on sequence homology of the two ORFs, varied between 2 to 633 nucleotides. Due to length differences and relative INDELs between the original ORFs, recombined ORFs differed in length. However, all recombinations occurred in frame and no premature stop codons were introduced, suggesting that these chimeric ORFs might yield functional proteins.

**Table 2:** Overview of all recombinations observed in 55 isolates obtained after laboratory evolution of strain IMS0408 under simulated lager fermentation conditions.

| Recombination<sup>1,2</sup> | Affected isolates | Locus 1 |  |  | Locus 2 |  |  | Homology<sup>5</sup> | Length chimeric ORF (bp) |
| --- | --- | --- | --- | --- | --- | --- | --- | --- | --- |
|  |  | Name<sup>1</sup> | Length<sup>3</sup> | Recombination<sup>4</sup> | Name<sup>1</sup> | Length<sup>3</sup> | Recombination<sup>4</sup> |  |  |
| ΔSe(YAR050W-YARend)<br>::Se(YALend-YAL063C) | IMS0614 | SeFLO1 | 5517 | 711-734 | SeFLO9 | 4752 | 712-735 | 79.23 % | 4752 |
| ΔSe::Sc(YBR275C-YBRend) | IMS0561 | ScRIF1 | 5751 | 562-572 | SeRIF1 | 5751 | 569-579 | 77.49 % | 5745 |
| ΔSc::Se(YDR051C-YDRend) | IMS0567 | SeDET1 | 1005 | 426-434 | ScDET1 | 1005 | 427-435 | 84.36 % | 1005 |
| ΔSc::Se(YGR218C-YGRend) | IMS0558 | ScCRM1 | 3251 | 2626-2636 | SeCRM1 | 3251 | 2627-2637 | 85.84 % | 3251 |
| ΔSc::Se(YGR271W-YGRend) | IMS0567 | ScSLH1 | 5904 | 1335-1349 | SeSLH1 | 5901 | 1336-1350 | 82.41 % | 5901 |
| ΔSc(YGR279C-YGRend)<br>::Se(YMR305C-YMRend) | IMS0563 | SeSCW10 | 1146 | 892-899 | ScSCW4 | 1161 | 908-915 | <70 %* | 1146 |
| ΔSc::Se(YGR282C-YGRend) | IMS0554 | SeBGL2 | 942 | 115-125 | ScBGL2 | 942 | 116-126 | 87.75 % | 942 |
| ΔSc::Se(YHLend-YHL023C) | IMS0552 | SeNPR3 | 3444 | 1551-1571 | ScNPR3 | 3441 | 1561-1581 | 78.80 % | 3432 |
| ΔSe::Sc(YKLend-YKL057C),<br>ΔSc::Se(YKL057C-YKrend) | IMS0548 | SeNUP120 | 3114 | 1867-1868 | ScNUP120 | 3114 | 1868-1869 | 80.14 % | 3114 |
| ΔSc::Se(YLR154C-YLRend) | IMS0544 | Ribosomal DNA | - | - | Ribosomal DNA | - | - | - | - |
| ΔSe::Sc(YLR154C-YLRend) | IMS0599 | Ribosomal DNA | - | - | Ribosomal DNA | - | - | - | - |
| ΔSc::Se(YLR305C-YLRend) | IMS0604 | SeSTT4 | 5703 | 5013-5018 | ScSTT4 | 5703 | 5014-5019 | 81.64 % | 5703 |
| ΔSc::Se(YNL061C-YNL055C) | IMS0562 | ScNOP2 | 1857 | 1386-1391 | SeNOP2 | 1860 | 1390-1395 | 87.11 % | 1857 |
|  |  | ScPOR1 | 852 | 399-401 | SePOR1 | 852 | 400-402 | 85.82 % | 852 |
| ΔSe::Sc(YNLend-YNL123W) | IMS0603 | ScNMA111 | 2994 | 303-314 | SeNMA111 | 2994 | 304-315 | 83.61 % | 2994 |
| ΔSc::Se(YOL013C-YOL006C) | IMS0603 | SeTOP1 | 2304 | 2010-2021 | ScTOP1 | 2311 | 2017-2028 | 82.64 % | 2304 |
| ΔSe::Sc(YOLend-YOL013C) | IMS0603 | SeHRD1 | 1644 | 312-329 | ScHRD1 | 1656 | 313-330 | 82.82 % | 1656 |
| ΔSe::Sc(YOLend-YOL057W) | IMS0597 | ScYOL057W | 3015 | 99-116 | SeYOL057W | 2133 | 100-117 | <70 %* | 2133 |
| ΔSe::Sc(YOLend-YOL072W) | IMS0595 | ScTHP1 | 1368 | 726-731 | SeTHP1 | 1374 | 733-738 | 79.07 % | 1368 |
| ΔSe::Sc(YOLend-YOL075W) | IMS0600 | SeYOL075C | 3903 | 2409-2414 | ScYOL075C | 3885 | 2392-2397 | 81.85 % | 3903 |
| ΔSe::Sc(YOR063W-YORend) | IMS0560 | ScRPL3 | 1164 | 1023-1063 | SeRPL3 | 1164 | 1024-1064 | 94.59 % | 1164 |
| ΔSe::Sc(YOR133W-YORend) | IMS0559, IMS0561 and IMS0562 | ScEFT1 | 2529 | 246-311 | SeEFT1 | 2529 | 247-312 | 94.94 % | 2529 |
| ΔSc(YCLend-YCL067C),<br>ΔSc(YCR039C-YCRend) | IMS0551 | ScHMLALPHA2 | 633 | 1-633 | ScMATALPHA2 | 633 | 1-633 | 100 % | 633 |
| ΔSc(YDR261C-YDR211W) | IMS0558 and IMS0560 | TY-transposon | - | - | TY-transposon | - | - | - | - |
| ΔSc(YKL032C-YKL054C) | IMS0543-547, IMS0599, IMS0601 and IMS0602 | ScIXR1 | 1794 | 944-955 | ScDEF1 | 2217 | 1281-1292 | <70 %* | 1881 |
| ΔSc(YKL032C-YKL054C) | IMS0604-608 | ScIXR1 | 1794 | 332-341 | ScDEF1 | 2217 | 1272-1281 | <70 %* | 1278 |

Footnotes: <sup>1</sup> Sc and Se indicate sequences on the *S. cerevisiae* and *S. eubayanus* subgenomes, respectively. <sup>2</sup> “ $\Delta$ ” indicates deletion and “::” indicates substitution. <sup>3</sup> Length of the affected ORF in bp. <sup>4</sup> Nucleotide coordinates of the locus at which recombination could have occurred. Due to the high homology of Locus 1 and Locus 2, they shared identical sequence stretches within which the recombination could have occurred. Recombination sites are therefore indicated by a range of nucleotide coordinates within the ORFs. <sup>5</sup> Homology between Locus 1 and Locus 2 as determined by sequence alignment.

### *IRA2*, *SFL1* and *MAL11* are mutated in multiple evolved isolates

A total of 76 SNPs and 43 INDELs were identified in the genomes of the 55 isolates (Supplementary Data File 1A and 1B), of which 38 SNPs and 17 INDELs occurred in ORFs and were non-synonymous (Table 1). Gene ontology analysis of all genes affected by non-synonymous SNPs or INDELs did not yield a significant enrichment in specific biological processes, molecular functions or cellular components. However, the genes *IRA2*, *SFL1* and *MAL11* were affected in more than one strain. *IRA2* encodes a RAS GTPase-activating protein, which is disrupted in many *S. cerevisiae* genomes from the CEN.PK strain family (Tanaka, et al. 1990; Tanaka, et al. 1991; Nijkamp, et al. 2012). In strain IMS0408, the *ScIRA2* was indeed disrupted while the *SeIRA2* ORF was intact. However, *SeIRA2* was mutated in 6/10 isolates from LG12.1 after 232 generations. *SeIRA2* had a frameshift in IMS0594, a premature stop codon in IMS0596 and was completely lost in four isolates due to different loss of heterozygosities: Δ*Se*::*Sc*(YOL_end_-YOL072W) in IMS0595, Δ*Se*::*Sc*(YOL_end_-YOL057W) in IMS0597, Δ*Se*::*Sc*(YOL_end_-YOL075W) in IMS0600 and ΔSe::Sc(YOL_end_-YOL013C) in IMS0603.

*SFL1* encodes a transcriptional repressor of flocculation genes, which was present both on *Sc*CHRXV and *Se*CHRVIII (Atsushi, et al. 1989). *ScSFL1* was mutated in 6/10 isolates from LG30.1 after 464 generations, which harboured a non-conservative substitution at the 605^th^ nucleotide, affecting its DNA binding domain (Atsushi, et al. 1989). *SeSFL1* had a frameshift in IMS0558 (LG30.1), a single nucleotide substitution in IMS0614 and IMS0617 (HG12.2) and was completely lost in four isolates of LG30.1 due to two losses of heterozygosity: Δ*Se*::*Sc*(YOR133W-YOR_end_) in IMS0559, IMS0561 and IMS0562, and Δ*Se*::*Sc*(YOR063W-YOR_end_) in IMS0560 (Table 1).

*ScMAL11*, also referred to as *AGT1*, encodes the only maltose transporter of the *MAL* gene family which enables efficient uptake of maltotriose in IMS0408 (Alves, et al. 2008). Sc*MAL11* is located on the right arm of *Sc*CHRVII and is absent in the *S. eubayanus* subgenome of IMS0408, which has no other maltotriose transporters (Hebly, et al. 2015; Brickwedde, et al. 2018). *ScMAL11* had a frameshift in IMS0557 (LG30.1) and lost its start codon in IMS0565 (LG30.2) (Table 1). In addition, *ScMAL11* was completely lost due to three different losses of heterozygosity: Δ*Sc*::*Se*(YGR282C-YGR_end_) in IMS0554 (LG30.1), Δ*Sc*::*Se*(YGR218C-YGR_end_) in IMS0558 (LG30.1) and Δ*Sc*::*Se*(YGR271W-YGR_end_) in IMS0567 (LG30.2), and due to the non-conservative recombination Δ*Sc*(YGR279C-YGR_end_)::*Se*(YMR305C-YMR_end_) in IMS0563 (LG30.2).

### Mutations in *SFL1* cause emergence of calcium-dependent flocculation

Eight of the 55 evolved isolates showed SNPs, INDELs or loss of heterozygosity in the flocculation inhibitor gene *SFL1*. In isolates IMS0558, IMS0559, IMS0560, IMS0561 and IMS0562 from LG30.1, both *SeSFL1* and *ScSFL1* were mutated, while in isolate IMS0556 from LG30.1 only *ScSFL1* was mutated and in isolates IMS0614 and IMS0617 from HG12.2, only *SeSFL1* was mutated. Evolved isolates that carried mutations in both *ScSFL1* and *SeSFL1* formed elevated conically-shaped colonies on YPD agar, while strain IMS0408 and evolved isolates with either an intact *SeSFL1* or *ScSFL1* did not (Figure 3A). Strains with mutations in both *ScSFL1* and *SeSFL1* also showed rapid sedimentation in micro-aerobic cultures on wort, which was not observed for the other evolved isolates or for strain IMS0408 (Figure 3F).

**Figure 3:**
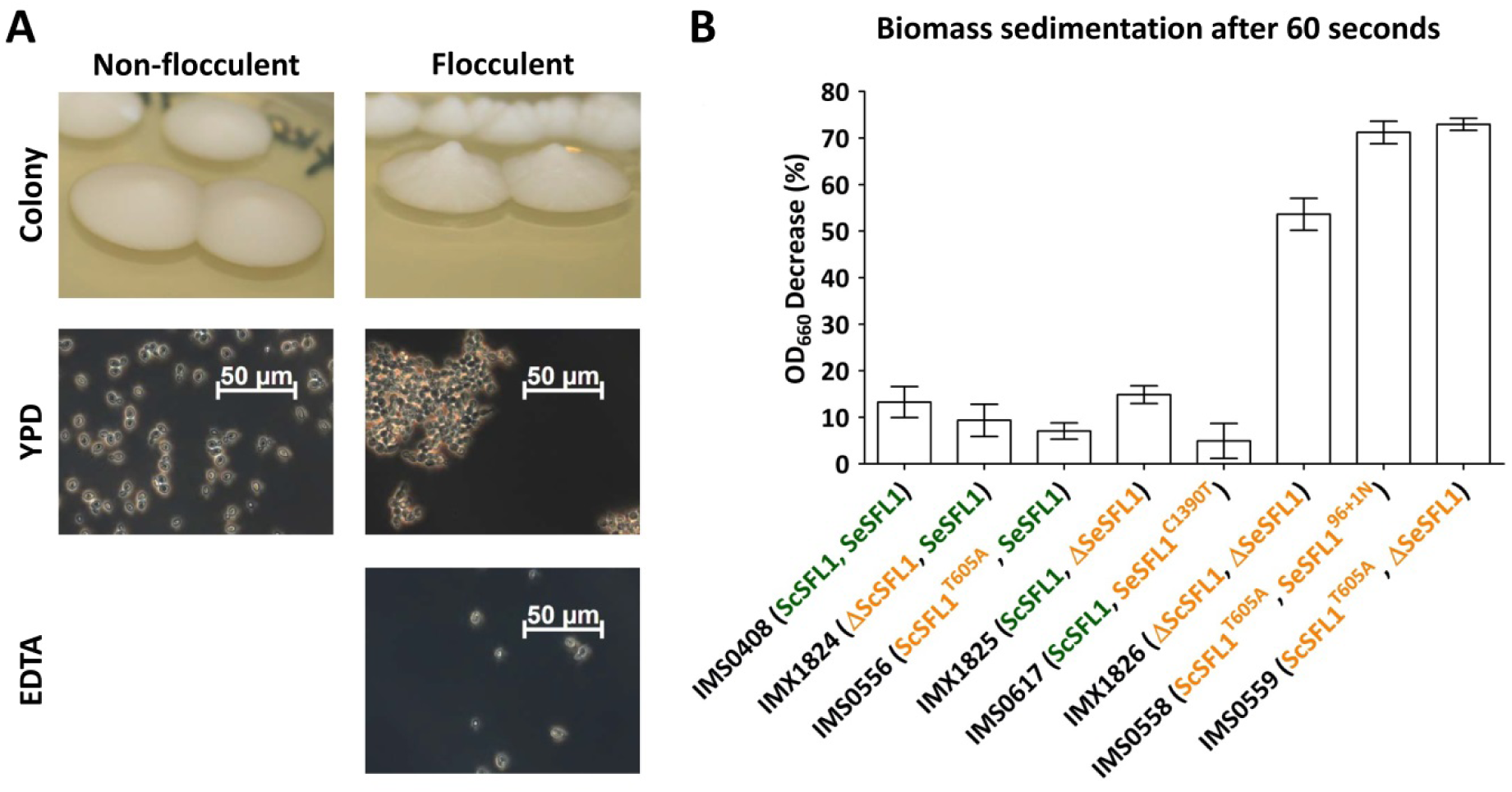
Mutations in *ScSFL1* and *SeSFL1*correlate with flocculation in evolved isolates and reverse engineered strains. A) Colony morphology and phase-contrast microscopy images (100x) of YPD-grown cell suspensions of the non-evolved, non-flocculent strain (IMS0408) and of a typical flocculent evolved isolate (IMS0558). Resuspension in 50 mM EDTA (pH 7.0) eliminated flocculation. B) Biomass sedimentation of evolved isolates and engineered strains with mutations in *SeSFL1* and/or *ScSFL1*. Triplicate cultures of all strains were grown on YPD and sedimentation was measured as the decrease in OD_660_ right underneath the meniscus of a stationary cell suspensions 60 s after the suspension had been vortexed.

*SFL1* is a repressor of *FLO* effector genes, which mediate calcium-dependent flocculation in *S. pastorianus* strains. To test if the acquired mutations in *SFL1* affected calcium-dependent flocculation, cultures from each strain were inspected microscopically. In the absence of EDTA, all sedimenting strains formed large aggregates, while IMS0408 and strains with an intact *SeSFL1* or *ScSFL1* did not (Figure 3A). Upon resuspension in Ca^2+^-chelating EDTA buffer, the aggregates reverted into a single-cell suspensions, thus confirming Ca^2+^-dependent flocculation. To test if the mutations affecting *SeSFL1* and *ScSFL1* were responsible for the calcium-dependent flocculation, single and double knockout strains were made using CRISPR-Cas9 gene editing: IMX1824 (Δ*ScSFL1*::mTurquoise2), IMX1825 (Δ*SeSFL1*::Venus) and IMX1826 (Δ*ScSFL1*::mTurquoise2, Δ*SeSFL1*::Venus). Flocculation intensity was assessed by measuring the decrease in OD_660_ at the surface of stationary-phase cultures after 1 minute without shaking. Isolates with mutations affecting only *SeSFL1* or only *ScSFL1* did not sediment significantly faster than IMS0408, which showed a decrease in OD_660_ of 13.3 ± 5.8 % (Figure 3B). However, IMX1826 (mTurquoise2::Δ*ScSFL1,* Venus::Δ*SeSFL1*), IMS0558 (*ScSFL1*^T605A^, *SeSFL1*^96+1N^) and IMS0559 (*ScSFL1*^T605A^, *ΔSeSFL1*) all showed decreases in OD_660_ above 53.6 %, indicating that disruption of both *ScSFL1* and *SeSFL1* causes a significant increase in flocculation (Student’s T test, p<0.0011). Since *SeSFL1* harboured three different mutations in isolates IMS0558, IMS0559, IMS0560, IMS0561 and IMS0562, calcium-dependent flocculation emerged by at least three independent mutations during the laboratory evolution experiment LG30.2.

### Mutations in *ScMAL11* cause loss of maltotriose utilisation

*ScMAL11*, which encodes the sole maltotriose transporter in strain IMS0408, was mutated in six evolved isolates. To investigate if these mutations affected maltotriose fermentation, the isolates were grown micro-aerobically on diluted wort at 30 °C. The unevolved IMS0408 consumed 100 ± 0 % of the maltotriose in diluted wort, and evolved strains with an intact *ScMAL11* genes consumed 98 ± 3 %. In contrast, strains IMS0554, IMS0557, IMS0558, IMS0563, IMS0565 and IMS0567, which all harboured mutations in *ScMAL11*, did not show any maltotriose consumption; instead, the concentration increased by 14 ± 3 % on average, presumably due to water evaporation. To test if the mutations affecting *ScMAL11* were responsible for the loss of maltotriose utilisation, *ScMAL11* was deleted in strain IMS0408 using CRISPR-Cas9 gene editing, resulting in strain IMX1698 (Δ*ScMAL11*::mVenus). Under the same conditions used to evaluate maltotriose utilization by the evolved strains, strain IMS0408 consumed 97 ± 5 % of the maltotriose while strain IMX1698 only consumed 1 ± 0 % of the maltotriose. These results confirmed that loss of *ScMAL11* function was responsible for loss of maltotriose utilization.

## Discussion

Evolution of the laboratory *S. cerevisiae* x *S. eubayanus* IMS0408 under simulated lager-brewing conditions yielded a wide array of mutations, including SNPs, INDELs, chromosomal recombinations, aneuploidy and loss of mitochondrial DNA (Table 1, Additional file 2). SNPs were the most common type of mutation, with frequencies ranging between 0.004 and 0.039 per division (Figure 4). Based on a genome size of 24.2 Mbp, the rate at which single-nucleotide mutations occurred was between 1.7∙10^−10^ and 1.6∙10^−9^ per nucleotide per cell division, which is similar to a rate of 3.3∙10^−10^ per site per cell division reported for *S. cerevisiae* (Lynch et al. 2008). At a frequency of 0.003 and 0.010 per cell division, INDELs occurred up to 4.1-fold less frequently than SNPs, in accordance with their twofold lower occurrence in *S. cerevisiae* (Lang and Murray 2008). The higher incidence of both SNPs and INDELs in isolates evolved in full-strength wort (Figure 4) may be related to the higher concentrations of ethanol, a known mutagen, in these cultures (Voordeckers, et al. 2015). The rate of loss of mitochondrial DNA varied between 0.0001 and 0.007 per division (Figure 4), and was negatively correlated with the number of generations of selective growth, indicating loss of mitochondrial DNA is selected against. The percentage of respiratory deficient isolates, between 13 and 40 % for all evolutions, is consistent with observations during laboratory evolution under oxygen limitation, and has been associated with increased ethanol tolerance (Ibeas and Jimenez 1997; Taylor, et al. 2002). However, loss of respirative capacity is highly undesirable for lager brewing as it impedes biomass propagation (Gibson, et al. 2008).

**Figure 4:**
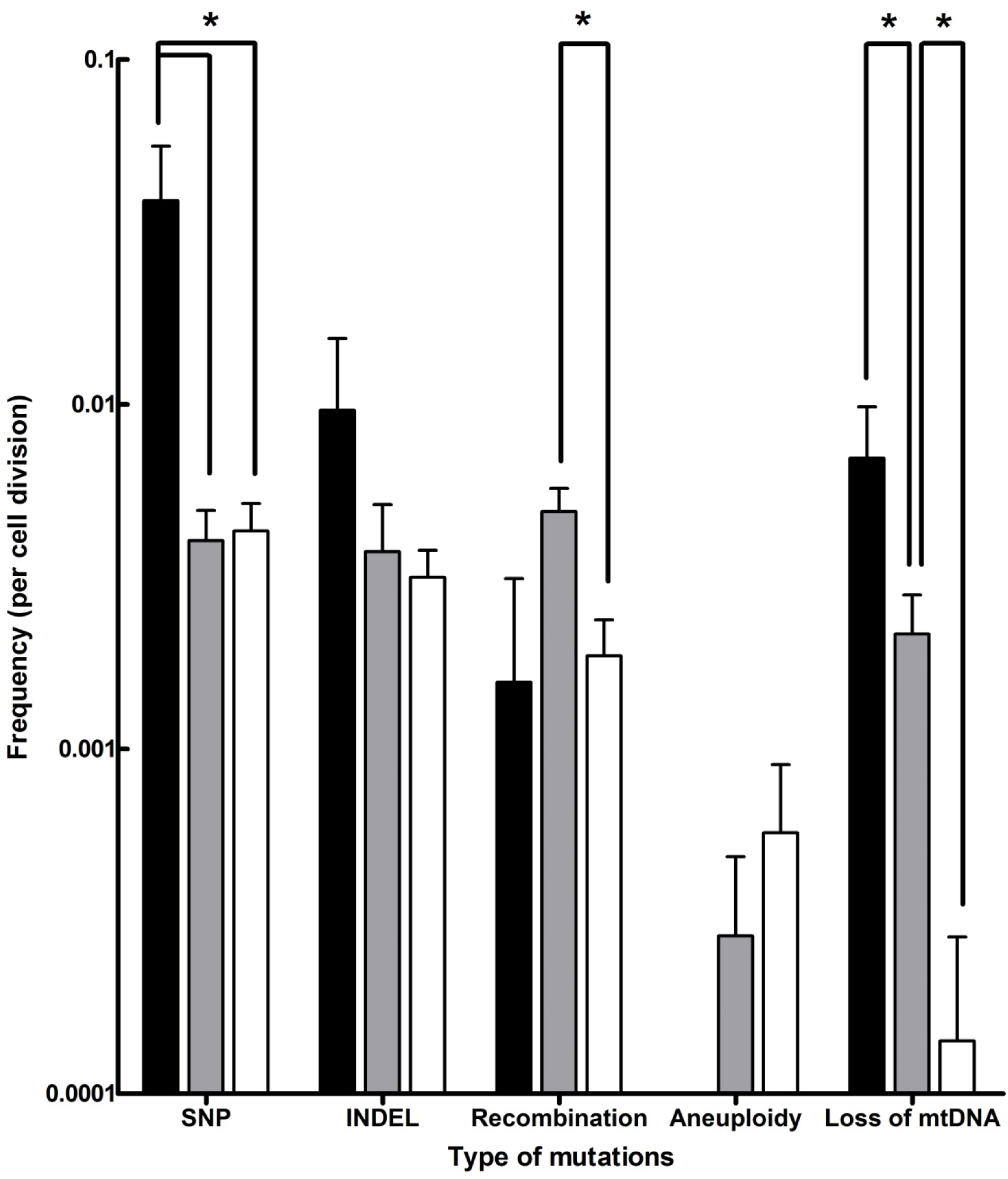
Frequencies of different types of mutations observed in evolved isolates obtained after laboratory evolution of strain IMS0408 under simulated lager fermentation conditions. The mutations identified in all 55 isolates evolved under brewing conditions were classified by type, and the frequency of mutation per cell division was calculated for each isolate based on its estimated number of generations of growth under simulated brewing conditions. Average frequencies of mutation types and standard deviations are shown for isolates evolved on full-strength wort at 12 °C (black), on three-fold diluted wort at 12 °C (grey) and on threefold diluted wort at 30 °C (white). The frequencies are shown on a logarithmic scale, and p-values were determined using Student’s t-test.

The frequency of chromosomal recombinations was estimated between 0.002 and 0.005 per division (Figure 4), which is similar to frequencies reported for *S. cerevisiae* (Dunham, et al. 2002). The observed recombinations were not reciprocal translocations. Instead, in all cases, genetic material was lost due to internal deletions, or genetic material from one chromosome was replaced by an additional copy of genetic material from another chromosome. This abundant loss of heterozygosity is consistent with the evolutionary history of *S. cerevisiae* (Magwene, et al. 2011) and with previously observed loss of genetic material in hybrids (Sipiczki 2008; Peris, et al. 2017; Smukowski Heil, et al. 2017; Smukowski Heil, et al. 2018). Under selective conditions, regions from one subgenome can be preferentially affected by loss of heterozygosity due to the fitness effects of genes they harbour. Due to its irreversibility, loss of heterozygosity is a determining mechanism for the evolution of hybrids during domestication (Pérez Través, et al. 2014). For example, in an *S. cerevisiae* x *S. uvarum* hybrid, the chromosomal region harbouring *ScPHO84* was preferentially retained at 30 ° C, while it was lost at the expense of its *S. uvarum* homolog at 15 ° C (Smukowski Heil, et al. 2017; Smukowski Heil, et al. 2018). Similarly, the superior growth of *S. cerevisiae* at 30 °C relative to *S. paradoxus* could be linked to *S. cerevisiae* alleles of 8 genes by investigating the effect of loss of heterozygosity in a laboratory hybrid (Weiss, et al. 2018). Moreover, the deletion of *S. cerevisiae* alleles in *S. uvarum* x *S. cerevisiae* had strong and varying impacts on fitness under glucose-, sulphate- and phosphate-limitation (Lancaster, et al. 2018). Overall, loss of heterozygosity is an irreversible process which enables rapid adaptation of hybrid genomes to the selective pressure of their growth environment. In the present study, loss of heterozygosity notably affected *ScMAL11* and *SeSFL1*, contributing to the acquired flocculation and loss of maltotriose utilization phenotypes. Other observed losses of heterozygosity may be due to genetic drift, or they may yield a selective advantage which we did not identify. Loss of heterozygosity may be advantageous by removing unfavourable dominant alleles, by enabling the expression of favourable recessive alleles, or by resolving redundancy, cross-talk and possible incompatibility between alleles and genes of both subgenomes (Piatkowska, et al. 2013; Gibson and Liti 2015).

Whole-chromosome aneuploidy was the rarest type of mutation, occurring only between 0 and 0.0003 times per division (Figure 4). The emergence of single chromosome aneuploidy in 9 % of the isolates after laboratory evolution is similar to observations during laboratory evolution of *S. cerevisiae* (Yona, et al. 2012; Voordeckers, et al. 2015; González-Ramos, et al. 2016; Gorter de Vries, Pronk, et al. 2017). However, the observed small extent of aneuploidy starkly contrasts with the massive aneuploidy of *S. pastorianus* brewing strains (Van den Broek, et al. 2015; Okuno, et al. 2016). These differences might of course be attributed to the difference in time scale between four months of laboratory evolution and several centuries of domestication. However, they might also be due to differences between the *S. cerevisiae* x *S. eubayanus* IMS0408 evolved in this study and the ancestral *S. pastorianus* hybrid. Firstly, IMS0408 was obtained by crossing a haploid *S. cerevisiae* strain with a haploid *S. eubayanus* spore (Hebly, et al. 2015). In contrast, the progenitor of brewing strains of *S. pastorianus* may have resulted from a cross of higher ploidy strains (Okuno, et al. 2016). Since higher ploidy leads to higher chromosome missegregation rates, they accumulate chromosome copy number changes faster (Storchova 2014). In addition, the higher initial ploidy leads to a smaller relative increase of genetic material when an additional copy is gained, which may ‘buffer’ deleterious effects of further changes in ploidy (Torres, et al. 2007). Moreover, the ancestral *S. pastorianus* hybrid may have had, or have acquired, mutations that stimulate extensive aneuploidy, such as mutations increasing the rate of chromosome missegregation or mutations increasing the tolerance against aneuploidy associated stresses (Torres, et al. 2010). Regardless of the origin of the extensive aneuploidy of *S. pastorianus*, our results show that euploid *S. cerevisiae* x *S. eubayanus* hybrids are not by definition prone to extensive aneuploidy under brewing-related experimental conditions. For industrial applications, the relative genetic stability of newly generated *Saccharomyces* hybrid strains reduces the chance of strain deterioration during the many generations involved in large-scale fermentation and/or biomass recycling (Krogerus, et al. 2017).

In addition to showing extensive aneuploidy, *S. pastorianus* strains harbour numerous chromosomal recombinations. During the laboratory evolution experiments, two types of recombinations were observed: (i) intrachromosomal recombinations resulting in loss of chromosome segments, and (ii) interchromosomal recombinations resulting in loss of one chromosome segment and replacement by an additional copy of a segment from another chromosome, resulting in loss of heterozygosity. While in *S. cerevisiae*, chromosomal recombinations predominantly occur in repetitive regions of the genome (Dunham, et al. 2002; Fontdevila 2005), here 88 % of the observed recombinations occurred within ORFs. The average homology of the recombined ORFs did not exceed the average 85 % homology of ORFs in the *S. cerevisiae* and *S. eubayanus* subgenomes. Instead, the high rate of recombinations at ORFs could reflect a correlation between transcriptional activity and recombination (Thomas and Rothstein 1989). In all cases, the reading frames were conserved, resulting in chimeric ORFs which could encode functional chimeric proteins, with altered length and sequences compared to the parental genomes. The potential selective advantage of such chimeric proteins is illustrated by recurring recombinations between ammonium permease *MEP2* alleles in a *S. cerevisiae* and *S. uvarum* hybrid during laboratory evolution under nitrogen limitation conditions (Dunn, et al. 2013). The formation of chimeric ORFs has even led to the emergence of novel gene functions, as illustrated by the formation of a maltotriose transporter by recombination of three non-maltotriose transporter genes in *S. eubayanus* during laboratory evolution (Brouwers, et al. 2018).

The predominant occurrence of recombinations within ORFs in the evolved isolates has also been observed in the genomes of brewing strains of *S. pastorianus*, which all share identical recombinations at the *ZUO1*, *MAT*, *HSP82* and *XRN1*/*KEM1* loci (Hewitt, et al. 2014; Walther, et al. 2014; Okuno, et al. 2016). These common recombinations suggest that all *S. pastorianus* isolates descend from a common ancestor (Monerawela and Bond 2018). However, since identical recombinations have been observed in independent evolutions, identical recombinations might reflect parallel evolution due to a strong selective advantage under brewing-related conditions and/or a predisposition of specific loci for recombination (Dunham, et al. 2002; Dunn, et al. 2013). Of the recombination loci found in the present study, only *EFT1* and *MAT* loci were associated with recombinations in *S. pastorianus.* Moreover, the recombinations at these loci in the evolved isolates were different from those in *S. pastorianus* (Hewitt, et al. 2014; Walther, et al. 2014; Okuno, et al. 2016; Monerawela and Bond 2017). All interchromosomal recombinations observed in this study were unique. These results, obtained under brewing-related conditions, are consistent with the notion that recombination sites are largely aleatory and that all modern *S. pastorianus* strains descend from a single hybrid ancestor.

Different recombination events resulted in the loss of heterozygosity, in four isolates each, of the right arm of *Se*CHRXV, including *SeSFL1*, and of the right arm of *Sc*CHRXI, including *ScMAL11*. These events directly affected two phenotypes relevant for brewing fermentation: calcium-dependent flocculation, which led to fast biomass sedimentation, and loss of maltotriose utilisation. Biomass sedimentation can be strongly selected for in sequential batch bioreactors, as it increases the chance that cells are retained in the bioreactor during the emptying phase (Oud, et al. 2013; Hope, et al. 2017). A similar selective advantage is likely to have played a role in the early domestication of *S. pastorianus,* as sedimenting yeast remaining in fermentation vessels was more likely to be used in a next fermentation. Flocculation is a key characteristic of current lager-brewing yeasts (also referred to as bottom-fermenting yeasts), as it simplifies biomass separation at the end of the fermentation (Ferreira, et al. 2010). The present study illustrates how this aspect of brewing yeast domestication can be rapidly reproduced under simulated laboratory conditions.

At first glance, loss of the ability to utilize maltotriose, an abundant fermentable sugar in wort, appears to be undesirable from an evolutionary perspective. However, as demonstrated in studies on laboratory evolution of *S. cerevisiae* in sequential batch cultures on sugar mixtures, the selective advantage of consuming a specific sugar from a sugar mixture correlates with the number of generations of growth on that sugar during each cultivation cycle (Wisselink, et al. 2009; Verhoeven, et al. 2018). *Saccharomyces* yeasts, including strain IMS0408 generally prefer glucose and maltose over maltotriose (Alves, et al. 2008; Hebly, et al. 2015; Brickwedde, et al. 2017). As a consequence, maltotriose consumption from wort typically only occurs when growth has already ceased due to oxygen limitation and/or nitrogen source depletion, which results in few or no generations of growth on this trisaccharide. However, loss of maltotriose utilization in six isolates in two independent evolution experiments strongly suggests that loss of *ScMAL11* expression was not merely neutral but even conferred a selective advantage. These results are consistent with the existence of many *S. pastorianus* strains with poor maltotriose utilisation and with the truncation of *ScMAL11* in all *S. pastorianus* strains, including good maltotriose utilizers (Vidgren, et al. 2009; Gibson, et al. 2013). In the latter strains, maltotriose utilisation depends on alternative transporters such as Mty1 (Dietvorst, et al. 2005; Salema-Oom, et al. 2005). It is therefore unclear if a selective advantage of the loss of *ScMAL11* reflects specific properties of this gene or its encoded transporter or, alternatively, a general negative impact of maltotriose utilisation under brewing-related conditions. In analogy with observations on maltose utilization by *S. cerevisiae*, unrestricted Mal11-mediated maltotriose-proton symport might cause maltotriose-accelerated death (Postma, et al. 1990; Jansen, et al. 2004). Alternatively, expression of the Mal11 transporter might compete with superior maltose transporters for intracellular trafficking, membrane integration and/or membrane space (Vidgren 2010; Libkind, et al. 2011). Indeed, laboratory evolution to obtain improved maltotriose utilisation resulted in reduced maltose uptake in *S. pastorianus* (Brickwedde, et al. 2017).

*Saccharomyces* hybrids are commonly applied for industrial applications such as lager beer brewing and wine fermentation (Naumov, et al. 2001; González, et al. 2006; Libkind, et al. 2011). Recently, novel hybrids have been generated, that performed well under a broad range of industrially-relevant conditions (Bellon, et al. 2011; Hebly, et al. 2015; Lopandic, et al. 2016; Krogerus, et al. 2017; Nikulin, et al. 2017; Peris, et al. 2017). While the performance and phenotypic diversity of laboratory hybrids support their application in industrial processes, further strain development of such hybrids could improve their performance (Steensels, et al. 2014; Krogerus, et al. 2017). Especially in food and beverage fermentation processes, consumer acceptance issues largely preclude use of targeted genetic modification techniques. Laboratory evolution offers an interesting alternative strategy for ‘non-GMO’ strain improvement (Bachmann, et al. 2015). However, as exemplified by the loss, in independent laboratory evolution experiments, of *MAL11* and of the mitochondrial genome, mutations that yield increased fitness under simulated industrial fermentation conditions are not necessarily advantageous for industrial performance. Therefore, instead of faithfully reconstructing industrial conditions in the laboratory, laboratory evolution experiments should be designed to specifically select for desired phenotypes. For example, a recent study illustrated how maltotriose fermentation kinetics of an *S. pastorianus* hybrid could be improved by laboratory evolution in carbon-limited chemostats grown on a maltotriose-enriched sugar mixture (Brickwedde, et al. 2017).

## Materials and Methods

### Yeast strains and media

*Saccharomyces* strains used in this study are listed in Supplementary Table 3. Yeast strains and *E. coli* strains containing plasmids were stocked in 1 mL aliquots after addition of 30 % v/v glycerol to the cultures and stored at −80 °C. For preparation of stock cultures and inocula of bioreactors, yeast strains were routinely propagated in shake flasks containing 100 mL YPD (10 g.L^−1^ yeast extract, 20 g.L^−1^ yeast peptone and 20 g.L^−1^ glucose) at 30 °C and 200 RPM in an Brunswick Innova43/43R shaker (Eppendorf Nederland B.V., Nijmegen, The Netherlands). For cultivation on solid media, YPD medium was supplemented with 20 g.L^−1^ Bacto agar (Becton Dickinson, Breda, The Netherlands) and incubation was done at 30 °C. Synthetic medium (SM), containing 3 g.L^−1^ KH_2_PO_4_, 0.5 g.L^−1^ MgSO_4_.7H2O, 5 g.L^−1^ (NH_4_)2SO_4_, 1 mL.L^−1^ of a trace element solution and 1 mL.L^−1^ of a vitamin solution, was prepared as previously described (Verduyn, et al. 1992). SM maltotriose was supplemented with 20 g.L^−1^ of maltotriose and SM ethanol with 20 mL.L^−1^ of ethanol. Selection for the amdS marker was performed on SM-AC: SM with 0.6 g∙L^−1^ acetamide and 6.6 g L^−1^ K_2_SO_4_ instead of (NH_4_)_2_SO_4_ as nitrogen source (Solis-Escalante, et al. 2013). For counter selection of the amdS marker, strains were first grown on YPD and then on SM-FAC: SM supplemented with 2.3 g∙L^−1^ fluoroacetamide (Solis-Escalante, et al. 2013). Industrial wort was provided by HEINEKEN Supply Chain B.V., Zoeterwoude, the Netherlands, and contained 14.4 g/L glucose, 2.3 g/L fructose, 85.9 g/L maltose, 26.8 g/L maltotriose and 269 mg/L free amino nitrogen. The wort was supplemented with 1.5 mg.L^−1^ Zn^2+^ by addition of ZnSO_4_.7H_2_O, then autoclaved for 30 min at 121 ᵒC and, prior to use, filtered through Nalgene 0.2 µm SFCA bottle-top filters (ThermoFisher Scientific, Waltham, MA). For experiments performed with diluted wort, two volumes of sterile demineralized water were added per volume of wort. To prevent excessive foaming during the aeration phase of the bioreactor experiments, (un)diluted wort was supplemented with 0.2 mL.L^−1^ of sterile Pluronic PE 6100 antifoam (Sigma-Aldrich, Zwijndrecht, the Netherlands).

**Table 3:** Plasmids used throughout this study.

| Name | Relevant genotype | Origin |
| --- | --- | --- |
| pUD711 | ori *bla* gRNA-*ScSFL1* | GeneArt™ |
| pUD712 | ori *bla* gRNA-*SeSFL1* | GeneArt™ |
| pUDE481 | ori *bla* ARS4/CEN6 *hyg*<sup>R</sup> *ScTDH3p*-mTurquoise2-*ScADH1t* | (Gorter de Vries, et al. 2018) |
| pUDE482 | ori *bla* ARS4/CEN6 *hyg*<sup>R</sup> *ScTEF1p*-Venus-*ScENO2t* | (Gorter de Vries, et al. 2018) |
| pUDP004 | ori *bla* panARSopt amdSYM *Spcas9* | (Gorter de Vries, de Groot, et al. 2017) |
| pUDP045 | ori *bla* panARSopt amdSYM *Spcas9* gRNA-*ScMAL11* | (Gorter de Vries, et al. 2018) |
| pUDP104 | ori *bla* panARSopt amdSYM *Spcas9* gRNA-*ScSFL1* | This study |
| pUDP105 | ori *bla* panARSopt amdSYM *Spcas9* gRNA-*SeSFL1* | This study |

### Analytical methods and statistics

Optical density at 660 nm was measured with a Libra S11 spectophotometer (Biochrom, Cambridge, UK). HPLC analysis of sugar and metabolite concentrations was performed with an Agilent Infinity 1260 chromatography system (Agilent Technologies, Santa Clara, CA) with an Aminex HPX-87 column (Bio-Rad, Lunteren, The Netherlands) at 65 °C, eluted with 5 mM H_2_SO_4_. Significance of data was assessed by an unpaired two-tailed Student’s t-test with a 95 % confidence interval.

### Laboratory evolution and single colony isolation

The hybrid yeast strain IMS0408 was evolved under three different conditions in duplicate in Minifors 2 bioreactors (INFORS HT, Velp, the Netherlands) with a working volume of 100 mL: on diluted wort at 30 °C (LG30.1 and LG30.2), on diluted wort at 12 °C (LG12.1 and LG12.2) and on full-strength wort at 12 °C (HG12.1 and HG12.2). Sequential batch cultivation was performed with 10 and 30 mL.min^−1^ of headspace N_2_ flushing at 12 and 30 °C, respectively. The percentage of CO_2_ in the outlet gas stream, the culture pH and the dissolved oxygen concentration in the broth were continuously monitored. The end of a batch cultivation cycle was automatically triggered when the percentage of CO_2_ in the offgas decreased below 75 % and 10 % of the maximum value reached during that cycle for growth on diluted wort and full-strength wort, respectively. These CO_2_ percentages correspond to the moment at which sugar utilisation was complete in the first batch cycle for each condition, as determined by HPLC measurements. When the CO_2_ threshold was reached, the reactor was emptied while stirring at 1200 RPM leaving about 7 mL to inoculate the next batch. Upon addition of fresh medium, the broth was stirred at 500 RPM and sparged with 500 mL.min^−1^ of pressurized air during 5 min for diluted wort or 12 h for wort. During the remainder of each batch cultivation cycle, the medium was not sparged or stirred and the pH was not adjusted. LG30.1 and LG30.2 were carried out for 116 and 117 cycles respectively, LG12.1 and LG12.2 were carried out for 29 cycles and HG12.1 and HG12.2 were carried out for 13 and 16 cycles, respectively. Culture samples from all six reactors were then streaked on YPD plates and after three subsequent restreaks, frozen stock cultures of single colony isolates were prepared. By default, five isolates were obtained for each culture. For LG12.1 and LG30.1, two different colony morphologies were observed, therefore five elevated and conically-shaped colonies and five regular flat colonies were stocked. The experiments at 12 °C on diluted wort were continued for four months until a total of 58 and 57 cycles was reached for LG12.1 and LG12.2, respectively, and five single-cell isolates were obtained for each reactor as described above.

### Genomic DNA extraction and whole genome sequencing

Yeast cultures were incubated in 500-mL shake-flasks containing 100 mL YPD at 30°C on an orbital shaker set at 200 RPM until the strains reached stationary phase at an OD_660_ between 12 and 20. Genomic DNA was isolated using the Qiagen 100/G kit (Qiagen, Hilden, Germany) according to the manufacturer’s instructions and quantified using a Qubit^®^ Fluorometer 2.0 (ThermoFisher Scientific). For IMS0408 and the evolved isolates, genomic DNA was sequenced at Novogene Bioinformatics Technology Co., Ltd (Yuen Long, Hong Kong) on a HiSeq2500 sequencer (Illumina, San Diego, CA) with 150 bp paired-end reads using PCR-free library preparation.

### Genome analysis

A high quality reference genome was constructed by combining near-complete assemblies of *S. cerevisiae* CEN.PK113-7D (Salazar, et al. 2017) and *S. eubayanus* CBS12357^T^ (Brickwedde, et al. 2018). The kanMX marker present in IMS0408 was inserted as an additional contig (Wach, et al. 1994). For each evolved strain, raw Illumina reads were aligned against the reference genome using the Burrows–Wheeler Alignment tool (BWA, version 0.7.15-r1142) and further processed using SAMtools (version 1.3.1) and Pilon (version 1.18) for variant calling (Li, et al. 2009; Li and Durbin 2010; Walker, et al. 2014). SNPs and INDELs that were also called or which were ambiguous in IMS0408, were disregarded. Copy number was determined based on read coverage analysis. Chromosomal translocations were detected using Breakdancer (version 1.3.6) (Chen, et al. 2009). Only translocations which were supported by at least 10 % of the reads aligned at that locus and which were absent in strain IMS0408 were considered. All SNPs, INDELs, recombinations and copy number changes were manually confirmed by visualising the generated .bam files in the Integrative Genomics Viewer (IGV) software (Robinson, et al. 2011). A complete list of identified mutations is provided in Supplementary Data File 1. For chimeric open-reading-frame reconstruction, reads aligning within 3 kbp of an identified recombination site and their paired reads were extracted using Python and were assembled using SPAdes (Bankevich, et al. 2012). The resulting contigs were aligned against ORFs of genes the genes affected by the recombination to identify the recombination point, and the complete recombined ORF was reconstructed. Original and recombined ORFs were then aligned and translated using CloneManager (version 9.51, Sci-Ed Software, Denver, CO) to determine whether the translocation had introduced frameshifts or premature stop codons.

### DNA content determination by flow cytometric analysis

Exponential-phase shake flask cultures on YPD were diluted to an OD_660_ of 1. A 1 mL sample (approximately 10^7^ cells) was then washed in cold demineralized water and resuspended in 800 μL 70 % ethanol while vortexing. After addition of another 800 μL 70 % ethanol, fixed cells were stored at 4 °C until further staining and analysis. DNA was then stained with SYTOX Green as described previously (Haase and Reed 2002). Samples were analysed on a Accuri C6 flow cytometer (BD Biosciences, Franklin Lakes, NJ) equipped with a 488-nm laser and the median fluorescence of cells in the 1n and 2n phases of the cell cycle was determined using FlowJo (BD Biosciences). The 1n and 2n medians of strains CEN.PK113-7D (n), CEN.PK122 (2n) and FRY153 (3n) were used to create a standard curve of fluorescence versus genome size with a linear curve fit, as performed previously (Van den Broek, et al. 2015). The genome size of each tested strain was estimated by averaging predicted genome sizes of the 1n and 2n population in assays on three independent cultures.

### Identification of strains with respiratory deficiency

Respiratory competence was assessed through their ability to grow on ethanol. Samples from 24 h shake-flask cultures on YPD (30 °C, 200 RPM) were washed twice with demineralized water and used to inoculate duplicate aerobic shake flasks containing 100 mL of SM with 2 % ethanol to an OD_660_ of 0.2. After 72 h incubation at 30 °C and 200 RPM, OD_660_ was measured.

### Assay for calcium-dependence of flocculation

Two 100 μL aliquots from overnight cultures on YPD were washed with sterile demineralized water. One aliquot was resuspended in demineralized water and the other in 50 mM EDTA (pH 7.0). Both samples were imaged at 100 x magnification under a Z1 microscope (Carl Zeiss BV, Breda, the Netherlands) to assess flocculence and its reversal by EDTA chelation of calcium ions.

### Plasmid construction

All plasmids were propagated in *E. coli* DH5α (Table 3). Plasmid pUD711 and pUD712 were designed as previously described (Gorter de Vries, de Groot, et al. 2017) and de novo synthesised at GeneArt (ThermoFisher Scientific) containing the sequence 5’ GGTCTCGCAAATAACAACTGATGAGTCCGTGAGGACGAAACGAGTAAGCTCGTCTTGTTATAGTCACGGATCGAGTTTTAGAGCTAGAAATAGCAAGTTAAAATAAGGCTAGTCCGTTATCAACTTGAAAAAGTGGCACCGAGTCGGTGCTTTTGGCCGGCATGGTCCCAGCCTCCTCGCTGGCGCCGGCTGGGCAACATGCTTCGGCATGGCGAATGGGACACAGGCAAAATTCATCTGATGAGTCCGTGAGGACGAAACGAGTAAGCTCGTCATGAATATCGCATTTTGTGGGTTTTAGAGCTAGAAATAGCAAGTTAAAATAAGGCTAGTCCGTTATCAACTTGAAAAAGTGGCACCGAGTCGGTGCTTTTGGCCGGCATGGTCCCAGCCTCCTCGCTGGCGCCGGCTGGGCAACATGCTTCGGCATGGCGAATGGGACACAGCGAGACC 3’ for pUD711 and 5’ GGTCTCGCAAATAACAACTGATGAGTCCGTGAGGACGAAACGAGTAAGCTCGTCTTGTTATAGTCACGGATCGAGTTTTAGAGCTAGAAATAGCAAGTTAAAATAAGGCTAGTCCGTTATCAACTTGAAAAAGTGGCACCGAGTCGGTGCTTTTGGCCGGCATGGTCCCAGCCTCCTCGCTGGCGCCGGCTGGGCAACATGCTTCGGCATGGCGAATGGGACACAGCGAGACC 3’ for pUD712. Plasmid pUDP104, expressing gRNA_*ScSFL1*_ and *cas9*, was constructed by Golden Gate cloning by digesting pUDP004 and pUD711 using BsaI and ligating with T4 ligase (Engler, et al. 2008). Similarly, plasmid pUDP105, expressing gRNA_*SeSFL1*_ and *cas9*, was constructed from pUDP004 and pUD712. Correct assembly was verified by restriction analysis using PdmI.

### Strain construction

The S*cTEF1*p-Venus-*ScENO2*t repair fragment with flanks for homologous recombination in the *ScMAL11* locus was PCR amplified from plasmid pUDE481 using primers 12989 and 12990 (Table 4). The *ScTDH3*p-mTurquoise2-*ScADH1*t repair fragment with flanks for homologous recombination in the *ScSFL1* locus was PCR amplified from plasmid pUDE482 using primers 13564 and 13565. The S*cTEF1*p-Venus-*ScENO2*t repair fragment with flanks for homologous recombination in the *SeSFL1* locus was PCR amplified from plasmid pUDE481 using primers 13566 and 13567.

**Table 4:**
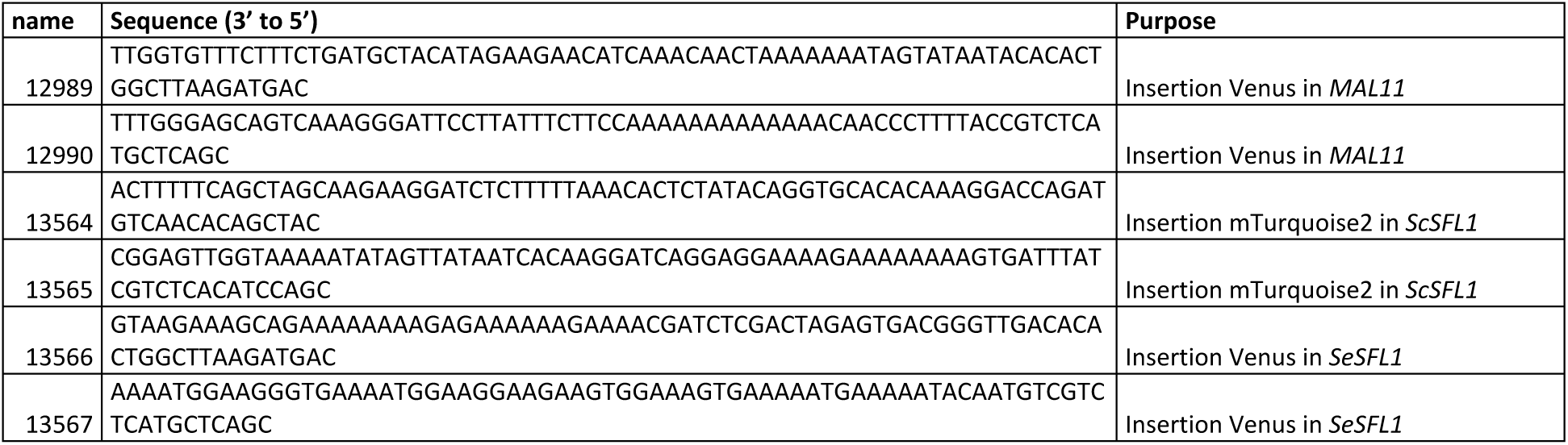
Primers used in this study.

All strains were transformed by electroporation as described previously, with 300 ng of gRNA/Cas9 expression plasmid and 1 μg of repair fragment (Gorter de Vries, de Groot, et al. 2017). Strains IMX1698 (mVenus::Δ*ScMAL11*), IMX1824 (mTurquoise2::Δ*ScSFL1*) and IMX1825 (Venus::Δ*SeSFL1*) were constructed by transforming IMS0408 with the appropriate repair fragments and plasmids pUDP045, pUDP104 and pUDP105, respectively. Strain IMX1826 (mTurquoise2::Δ*ScSFL1* Venus::Δ*SeSFL1*) was constructed by transforming IMX1824 using the appropriate repair fragment and plasmid pUDP105. After electroporation, cells were transferred to 20 mL SM-Ac medium to select successful transformants and incubated at 30 °C for 3 to 5 days. After growth was observed, 200 μL of culture was transferred to 20 mL fresh SM-Ac and incubated similarly during 24h. Finally, 200 μL from the second culture was transferred to 20 mL fresh YPD medium to maximize expression of fluorescent proteins. Successfully gene-edited cells were sorted using the BD FACSAria™ II SORP Cell Sorter (BD Biosciences) as described previously (Gorter de Vries, et al. 2018). The plasmids were cured from strains IMX1698, IMX1824, IMX1825 and IMX1826 by subsequent growth on YPD and plating on SM-FAC. After confirmation of the correct genotype by colony PCR, randomly picked colonies were used to prepare frozen stocks.

### Biomass sedimentation assay

IMS0408, IMS0556, IMS0558, IMS0559, IMS0617, IMX1824, IMX1825 and IMX1826 were grown in triplicate during 72h in vented 50 mL Bio-One Cellstar Cellreactor tubes (Sigma-Aldrich) on 20 mL YPD at 30 °C and 200 RPM until stationary phase. For each sample, the biomass was resuspended by vigorous vortexing and 1 mL was sampled immediately after vortexing from right underneath the meniscus. After 60 s of stationary incubation, another sample was taken by the same procedure. Biomass sedimentation was quantified as the ratio of the OD_660_ values of the two samples.

### Evaluation of maltotriose fermentation

Each strain was grown microaerobically in 100 mL serum bottles containing 100 mL medium and shaken at 200 RPM. Medium (full-strength or diluted wort) and incubation temperature (12 °C or 30 °C) were the same as in the evolution experiment from which a strain had been isolated. Strain IMS0408 was included as a control for each condition. Bottles were inoculated to an OD_660_ of 0.2 from aerobic shake-flask precultures grown on same medium and under the same conditions. During cultivation, 8 to 12 samples were taken at regular intervals for OD_660_ measurements and metabolite analysis by HPLC. When no further sugar consumption was recorded over an interval of at least 48h, the fermentation was considered finished.

## Acknowledgments and funding information

The sequencing data are available at NCBI (https://www.ncbi.nlm.nih.gov/) under the Bioproject PRJNA506072. We thank Erik de Hulster, Xavier Hakkaart and Robert Mans for sharing their expertise in fermentation, Marijke Luttik for her expertise in Flow cytometry and Nikola Gyurchev for assembling plasmids pUDP104 and pUDP105. We are thankful to Niels Kuijpers (Heineken Supply Chain B.V.), and Jan-Maarten Geertman (Heineken Supply Chain B.V.) for their support and for critically reading the manuscript. This work was performed within the BE-Basic R&D Program (http://www.be-basic.org/), which was granted an FES subsidy from the Dutch Ministry of Economic Affairs, Agriculture and Innovation (EL&I).

## Author contributions

ARGdV planned and executed the laboratory evolution, made single colony isolates, prepared genomic DNA and characterised ploidy and respiratory deficiency. ARGdV, ACAvA and LHK characterised flocculation and maltotriose utilisation of the isolates. ARGdV and MAV performed the reverse engineering of deletions of *SFL1* and *MAL11* and characterized the reverse engineered phenotypes. ARGdV and MvdB performed the bioinformatics analysis. ARGdV, JTP and JMGD wrote the manuscript. AS, NB and TA provided critical feedback throughout the study. JMGD and JTP supervised the study. All authors read and approved the final manuscript.

## References

Alves SL, Herberts RA, Hollatz C, Trichez D, Miletti LC, De Araujo PS, Stambuk BU. 2008. Molecular analysis of maltotriose active transport and fermentation by Saccharomyces cerevisiae reveals a determinant role for the AGT1 permease. Appl. Environ. Microbiol. 74:1494–1501.

Andreasen AA, Stier T. 1953. Anaerobic nutrition of Saccharomyces cerevisiae. I. Ergosterol requirement for growth in a defined medium. J Cell. Comp. Physiol. 41:23–36.

Atsushi F, Yoshiko K, Satoru K, Yoshio M, Shinichi M, Harumi K. 1989. Domains of the SFL1 protein of yeasts are homologous to Myc oncoproteins or yeast heat-shock transcription factor. Gene 85:321–328.

Bachmann H, Pronk JT, Kleerebezem M, Teusink B. 2015. Evolutionary engineering to enhance starter culture performance in food fermentations. Curr. Opin. Biotechnol. 32:1–7.

Bachmann H, Starrenburg MJ, Molenaar D, Kleerebezem M, van Hylckama Vlieg JE. 2012. Microbial domestication signatures of Lactococcus lactis can be reproduced by experimental evolution. Genome Res. 22:115–124.

Bankevich A, Nurk S, Antipov D, Gurevich AA, Dvorkin M, Kulikov AS, Lesin VM, Nikolenko SI, Pham S, Prjibelski AD. 2012. SPAdes: a new genome assembly algorithm and its applications to single-cell sequencing. J. Comput. Biol. 19:455–477.

Bellon JR, Eglinton JM, Siebert TE, Pollnitz AP, Rose L, de Barros Lopes M, Chambers PJ. 2011. Newly generated interspecific wine yeast hybrids introduce flavour and aroma diversity to wines. Appl. Microbiol. Biotechnol. 91:603–612.

Botstein D, Chervitz SA, Cherry M. 1997. Yeast as a model organism. Science 277:1259–1260.

Brickwedde A, Brouwers N, van den Broek M, Gallego Murillo JS, Fraiture JL, Pronk JT, Daran J-MG. 2018. Structural, physiological and regulatory analysis of maltose transporter genes in Saccharomyces eubayanus CBS 12357T. Front. Microbiol. 9:1786.

Brickwedde A, van den Broek M, Geertman J-MA, Magalhães F, Kuijpers NGA, Gibson B, Pronk JT, Daran J-MG. 2017. Evolutionary engineering in chemostat cultures for improved maltotriose fermentation kinetics in Saccharomyces pastorianus lager brewing yeast. Front. Microbiol. 8:1690.

Briggs DE, Brookes P, Stevens R, Boulton C. 2004. Brewing: science and practice: Elsevier.

Brouwers N, Gorter de Vries AR, van den Broek M, Weening SM, Schuurman TDE, Kuijpers NGA, Pronk JT, Daran J-MG. 2018. In vivo recombination of Saccharomyces eubayanus maltose-transporter genes yields a chimeric transporter that enables maltotriose fermentation. bioRxiv:428839.

Chambers SR, Hunter N, Louis EJ, Borts RH. 1996. The mismatch repair system reduces meiotic homeologous recombination and stimulates recombination-dependent chromosome loss. Mol. Cell Biol. 16:6110–6120.

Chen K, Wallis JW, McLellan MD, Larson DE, Kalicki JM, Pohl CS, McGrath SD, Wendl MC, Zhang Q, Locke DP. 2009. BreakDancer: an algorithm for high-resolution mapping of genomic structural variation. Nat. Methods 6:677.

Coloretti F, Zambonelli C, Tini V. 2006. Characterization of flocculent Saccharomyces interspecific hybrids for the production of sparkling wines. Food Microbiol. 23:672–676.

Delneri D, Colson I, Grammenoudi S, Roberts IN, Louis EJ, Oliver SG. 2003. Engineering evolution to study speciation in yeasts. Nature 422:68.

Dietvorst J, Londesborough J, Steensma H. 2005. Maltotriose utilization in lager yeast strains: MTT1 encodes a maltotriose transporter. Yeast 22:775–788.

Dunham MJ, Badrane H, Ferea T, Adams J, Brown PO, Rosenzweig F, Botstein D. 2002. Characteristic genome rearrangements in experimental evolution of Saccharomyces cerevisiae. Proc. Natl. Acad. Sci. U S A 99:16144–16149.

Dunn B, Paulish T, Stanbery A, Piotrowski J, Koniges G, Kroll E, Louis EJ, Liti G, Sherlock G, Rosenzweig F. 2013. Recurrent rearrangement during adaptive evolution in an interspecific yeast hybrid suggests a model for rapid introgression. PLoS Genet. 9:e1003366.

Engler C, Kandzia R, Marillonnet S. 2008. A one pot, one step, precision cloning method with high throughput capability. PLoS One 3:e3647.

Ferreira I, Pinho O, Vieira E, Tavarela J. 2010. Brewer’s Saccharomyces yeast biomass: characteristics and potential applications. Trends Food Sci. Technol. 21:77–84.

Fontdevila A. 2005. Hybrid genome evolution by transposition. Cytogenet. Genome Res. 110:49–55.

Gibbons JG, Rinker DC. 2015. The genomics of microbial domestication in the fermented food environment. Curr. Opin. Genet. Dev. 35:1–8.

Gibbons JG, Salichos L, Slot JC, Rinker DC, McGary KL, King JG, Klich MA, Tabb DL, McDonald WH, Rokas A. 2012. The evolutionary imprint of domestication on genome variation and function of the filamentous fungus Aspergillus oryzae. Curr. Biol. 22:1403–1409.

Gibson B, Liti G. 2015. Saccharomyces pastorianus: genomic insights inspiring innovation for industry. Yeast 32:17–27.

Gibson B, Prescott K, Smart K. 2008. Petite mutation in aged and oxidatively stressed ale and lager brewing yeast. Lett. Appl. Microbiol. 46:636–642.

Gibson BR, Lawrence SJ, Leclaire JP, Powell CD, Smart KA. 2007. Yeast responses to stresses associated with industrial brewery handling. FEMS Microbiol. Lett. 31:535–569.

Gibson BR, Storgårds E, Krogerus K, Vidgren V. 2013. Comparative physiology and fermentation performance of Saaz and Frohberg lager yeast strains and the parental species Saccharomyces eubayanus. Yeast 30:255–266.

González-Ramos D, Gorter de Vries AR, Grijseels SS, Berkum MC, Swinnen S, Broek M, Nevoigt E, Daran J-MG, Pronk JT, Maris AJA. 2016. A new laboratory evolution approach to select for constitutive acetic acid tolerance in Saccharomyces cerevisiae and identification of causal mutations. Biotechnol. Biofuels 9:173.

González SS, Barrio E, Gafner J, Querol A. 2006. Natural hybrids from Saccharomyces cerevisiae, Saccharomyces bayanus and Saccharomyces kudriavzevii in wine fermentations. FEMS Yeast Res. 6:1221–1234.

Gorter de Vries AR, Couwenberg LGF, van den Broek M, de la Torre Cortes P, ter Horst J, Pronk JT, Daran J-MG. 2018. Allele-specific genome editing using CRISPR-Cas9 causes off-target mutations in diploid yeast. bioRxiv:397984.

Gorter de Vries AR, de Groot PA, van den Broek M, Daran J-MG. 2017. CRISPR-Cas9 mediated gene deletions in lager yeast Saccharomyces pastorianus. Microb. Cell Fact. 16:222.

Gorter de Vries AR, Pronk JT, Daran J-MG. 2017. Industrial relevance of chromosomal copy number variation in Saccharomyces yeasts. Appl. Environ. Microbiol. 83:e03206–03216.

Haase SB, Reed SI. 2002. Improved flow cytometric analysis of the budding yeast cell cycle. Cell cycle 1:117–121.

Hebly M, Brickwedde A, Bolat I, Driessen MRM, de Hulster EAF, van den Broek M, Pronk JT, Geertman J-M, Daran J-MG, Daran-Lapujade P. 2015. S. cerevisiae × S. eubayanus interspecific hybrid, the best of both worlds and beyond. FEMS Yeast Res. 15:fov005.

Hewitt SK, Donaldson IJ, Lovell SC, Delneri D. 2014. Sequencing and characterisation of rearrangements in three S. pastorianus strains reveals the presence of chimeric genes and gives evidence of breakpoint reuse. PLoS One 9:e92203.

Hittinger CT. 2013. Saccharomyces diversity and evolution: a budding model genus. Trends Genet. 29:309–317.

Hope EA, Amorosi CJ, Miller AW, Dang K, Heil CS, Dunham MJ. 2017. Experimental evolution reveals favored adaptive routes to cell aggregation in yeast. Genetics:genetics. 116.198895.

Ibeas JI, Jimenez J. 1997. Mitochondrial DNA loss caused by ethanol in Saccharomyces flor yeasts. Appl. Environ. Microbiol. 63:7–12.

Jansen ML, Daran-Lapujade P, de Winde JH, Piper MD, Pronk JT. 2004. Prolonged maltose-limited cultivation of Saccharomyces cerevisiae selects for cells with improved maltose affinity and hypersensitivity. Appl. Environ. Microbiol. 70:1956–1963.

Krogerus K, Magalhães F, Vidgren V, Gibson B. 2017. Novel brewing yeast hybrids: creation and application. Appl. Microbiol. Biotechnol. 101:65–78.

Lancaster SM, Payen C, Heil CS, Dunham MJ. 2018. Fitness benefits of loss of heterozygosity in Saccharomyces hybrids. bioRxiv:452748.

Lang GI, Murray AW. 2008. Estimating the per-base-pair mutation rate in the yeast Saccharomyces cerevisiae. Genetics 178:67–82.

Li H, Durbin R. 2010. Fast and accurate long-read alignment with Burrows–Wheeler transform. Bioinformatics 26:589–595.

Li H, Handsaker B, Wysoker A, Fennell T, Ruan J, Homer N, Marth G, Abecasis G, Durbin R. 2009. The sequence alignment/map format and SAMtools. Bioinformatics 25:2078–2079.

Libkind D, Hittinger CT, Valério E, Gonçalves C, Dover J, Johnston M, Gonçalves P, Sampaio JP. 2011. Microbe domestication and the identification of the wild genetic stock of lager-brewing yeast. Proc. Natl. Acad. Sci. U S A 108:14539–14544.

Liti G, Barton DB, Louis EJ. 2006. Sequence diversity, reproductive isolation and species concepts in Saccharomyces. Genetics 174:839–850.

Lopandic K, Pfliegler WP, Tiefenbrunner W, Gangl H, Sipiczki M, Sterflinger K. 2016. Genotypic and phenotypic evolution of yeast interspecies hybrids during high-sugar fermentation. Appl. Microbiol. Biotechnol. 100:6331–6343.

Magwene PM, Kayikçi Ö, Granek JA, Reininga JM, Scholl Z, Murray D. 2011. Outcrossing, mitotic recombination, and life-history trade-offs shape genome evolution in Saccharomyces cerevisiae. Proc. Natl. Acad. Sci. U S A 108:1987–1992.

Marsit S, Dequin S. 2015. Diversity and adaptive evolution of Saccharomyces wine yeast: a review. FEMS Yeast Res. 15:fov067–fov067.

Meussdoerffer FG. 2009. A comprehensive history of beer brewing. Handbook of brewing: Processes, technology, markets:1–42.

Monerawela C, Bond U. 2018. The hybrid genomes of Saccharomyces pastorianus: A current perspective. Yeast 35:39–50.

Monerawela C, Bond U. 2017. Recombination sites on hybrid chromosomes in Saccharomyces pastorianus share common sequence motifs and define a complex evolutionary relationship between group I and II lager yeasts. FEMS Yeast Res. 17:fox047–fox047.

Naseeb S, James SA, Alsammar H, Michaels CJ, Gini B, Nueno-Palop C, Bond CJ, McGhie H, Roberts IN, Delneri D. 2017. Saccharomyces jurei sp nov., isolation and genetic identification of a novel yeast species from Quercus robur. Int. J. Syst. Evol. Microbiol. 67:2046–2052.

Naumov G, Nguyen H-V, Naumova ES, Michel A, Aigle M, Gaillardin C. 2001. Genetic identification of Saccharomyces bayanus var. uvarum, a cider-fermenting yeast. Int. J. Food Microbiol. 65:163–171.

Newlon C, Lipchitz L, Collins I, Deshpande A, Devenish R, Green R, Klein H, Palzkill T, Ren R, Synn S. 1991. Analysis of a circular derivative of Saccharomyces cerevisiae chromosome III: a physical map and identification and location of ARS elements. Genetics 129:343–357.

Nijkamp JF, van den Broek M, Datema E, de Kok S, Bosman L, Luttik MA, Daran-Lapujade P, Vongsangnak W, Nielsen J, Heijne WH. 2012. De novo sequencing, assembly and analysis of the genome of the laboratory strain Saccharomyces cerevisiae CEN. PK113-7D, a model for modern industrial biotechnology. Microb. Cell Fact. 11:36.

Nikulin J, Krogerus K, Gibson B. 2017. Alternative Saccharomyces interspecies hybrid combinations and their potential for low-temperature wort fermentation. Yeast.

Okuno M, Kajitani R, Ryusui R, Morimoto H, Kodama Y, Itoh T. 2016. Next-generation sequencing analysis of lager brewing yeast strains reveals the evolutionary history of interspecies hybridization. DNA Res. 23:67–80.

Oud B, Guadalupe-Medina V, Nijkamp JF, de Ridder D, Pronk JT, van Maris AJ, Daran J-M. 2013. Genome duplication and mutations in ACE2 cause multicellular, fast-sedimenting phenotypes in evolved Saccharomyces cerevisiae. Proc. Natl. Acad. Sci. U S A:201305949.

Pérez Través L, Lopes CA, Barrio Esparducer E, Querol Simón A. 2014. Stabilization process in Saccharomyces intra and interspecific hybrids in fermentative conditions. Int. Microbiol.

Peris D, Moriarty RV, Alexander WG, Baker E, Sylvester K, Sardi M, Langdon QK, Libkind D, Wang Q-M, Bai F-Y. 2017. Hybridization and adaptive evolution of diverse Saccharomyces species for cellulosic biofuel production. Biotechnol. Biofuels 10:78.

Piatkowska EM, Naseeb S, Knight D, Delneri D. 2013. Chimeric protein complexes in hybrid species generate novel phenotypes. PLoS Genet. 9:e1003836.

Postma E, Verduyn C, Kuiper A, Scheffers WA, Van Dijken JP. 1990. Substrate-accelerated death of Saccharomyces cerevisiae CBS 8066 under maltose stress. Yeast 6:149–158.

Querol A, Bond U. 2009. The complex and dynamic genomes of industrial yeasts. FEMS Microbiol. Lett. 293:1–10.

Robinson JT, Thorvaldsdóttir H, Winckler W, Guttman M, Lander ES, Getz G, Mesirov JP. 2011. Integrative genomics viewer. Nat. Biotechnol. 29:24.

Salazar AN, Gorter de Vries AR, van den Broek M, Wijsman M, de la Torre Cortés P, Brickwedde A, Brouwers N, Daran J-MG, Abeel T. 2017. Nanopore sequencing enables near-complete de novo assembly of Saccharomyces cerevisiae reference strain CEN. PK113-7D. FEMS Yeast Res. 17.

Salema-Oom M, Pinto VV, Gonçalves P, Spencer-Martins I. 2005. Maltotriose utilization by industrial Saccharomyces strains: characterization of a new member of the α-glucoside transporter family. Appl. Environ. Microbiol. 71:5044–5049.

Santaguida S, Amon A. 2015. Short-and long-term effects of chromosome mis-segregation and aneuploidy. Nat. Rev. Mol. Cell. Biol. 16:473.

Sheltzer JM, Blank HM, Pfau SJ, Tange Y, George BM, Humpton TJ, Brito IL, Hiraoka Y, Niwa O, Amon A. 2011. Aneuploidy drives genomic instability in yeast. Science 333:1026–1030.

Sipiczki M. 2008. Interspecies hybridization and recombination in Saccharomyces wine yeasts. FEMS Yeast Res. 8:996–1007.

Smukowski Heil CS, DeSevo CG, Pai DA, Tucker CM, Hoang ML, Dunham MJ. 2017. Loss of heterozygosity drives adaptation in hybrid yeast. Mol. Biol. Evol. 34:1596–1612.

Smukowski Heil CS, Large CR, Patterson K, Dunham MJ. 2018. Temperature preference biases parental genome retention during hybrid evolution. bioRxiv:429803.

Solis-Escalante D, Kuijpers NGA, Nadine B, Bolat I, Bosman L, Pronk JT, Daran J-MG, Pascale D-L. 2013. amdSYM, a new dominant recyclable marker cassette for Saccharomyces cerevisiae. FEMS Yeast Res. 13:126–139.

Steensels J, Snoek T, Meersman E, Nicolino MP, Voordeckers K, Verstrepen KJ. 2014. Improving industrial yeast strains: exploiting natural and artificial diversity. FEMS microbiology reviews 38:947–995.

Storchova Z. 2014. Ploidy changes and genome stability in yeast. Yeast 31:421–430.

Tanaka K, Lin BK, Wood DR, Tamanoi F. 1991. IRA2, an upstream negative regulator of RAS in yeast, is a RAS GTPase-activating protein. Proc. Natl. Acad. Sci. U S A 88:468–472.

Tanaka K, Nakafuku M, Satoh T, Marshall MS, Gibbs JB, Matsumoto K, Kaziro Y, Toh-e A. 1990. S. cerevisiae genes IRA1 and IRA2 encode proteins that may be functionally equivalent to mammalian ras GTPase activating protein. Cell 60:803–807.

Taylor DR, Zeyl C, Cooke E. 2002. Conflicting levels of selection in the accumulation of mitochondrial defects in Saccharomyces cerevisiae. Proc. Natl. Acad. Sci. U S A 99:3690–3694.

Thomas BJ, Rothstein R. 1989. Elevated recombination rates in transcriptionally active DNA. Cell 56:619–630.

Torres EM, Dephoure N, Panneerselvam A, Tucker CM, Whittaker CA, Gygi SP, Dunham MJ, Amon A. 2010. Identification of aneuploidy-tolerating mutations. Cell 143:71–83.

Torres EM, Sokolsky T, Tucker CM, Chan LY, Boselli M, Dunham MJ, Amon A. 2007. Effects of aneuploidy on cellular physiology and cell division in haploid yeast. Science 317:916–924.

Van den Broek M, Bolat I, Nijkamp JF, Ramos E, Luttik MAH, Koopman F, Geertman J-M, De Ridder D, Pronk J, Daran J-MG. 2015. Chromosomal copy number variation in Saccharomyces pastorianus is evidence for extensive genome dynamics in industrial lager brewing strains. Appl. Environ. Microbiol. 81:6253–6267.

Verduyn C, Postma E, Scheffers WA, Van Dijken JP. 1992. Effect of benzoic acid on metabolic fluxes in yeasts: a continuous-culture study on the regulation of respiration and alcoholic fermentation. Yeast 8:501–517.

Verhoeven MD, de Valk SC, Daran J-MG, van Maris AJ, Pronk JT. 2018. Fermentation of glucose-xylose-arabinose mixtures by a synthetic consortium of single-sugar-fermenting Saccharomyces cerevisiae strains. FEMS Yeast Res. 18:foy075.

Vidgren V. 2010. Maltose and maltotriose transport into ale and lager brewer’s yeast strains. VTT publications.

Vidgren V, Huuskonen A, Virtanen H, Ruohonen L, Londesborough J. 2009. Improved fermentation performance of a lager yeast after repair of its AGT1 maltose and maltotriose transporter genes. Appl. Environ. Microbiol. 75:2333–2345.

Voordeckers K, Kominek J, Das A, Espinosa-Cantú A, De Maeyer D, Arslan A, Van Pee M, van der Zande E, Meert W, Yang Y. 2015. Adaptation to high ethanol reveals complex evolutionary pathways. PLoS Genet. 11:e1005635.

Wach A, Brachat A, Pöhlmann R, Philippsen P. 1994. New heterologous modules for classical or PCR-based gene disruptions in Saccharomyces cerevisiae. Yeast 10:1793–1808.

Walker BJ, Abeel T, Shea T, Priest M, Abouelliel A, Sakthikumar S, Cuomo CA, Zeng Q, Wortman J, Young SK. 2014. Pilon: an integrated tool for comprehensive microbial variant detection and genome assembly improvement. PLoS One 9:e112963.

Walther A, Hesselbart A, Wendland J. 2014. Genome sequence of Saccharomyces carlsbergensis, the world’s first pure culture lager yeast. G3 (Bethesda) 4:783–793.

Weiss CV, Roop JI, Hackley RK, Chuong JN, Grigoriev IV, Arkin AP, Skerker JM, Brem RB. 2018. Genetic dissection of interspecific differences in yeast thermotolerance. Nat. Genet.:1.

Wisselink HW, Toirkens MJ, Wu Q, Pronk JT, van Maris AJA. 2009. Novel evolutionary engineering approach for accelerated utilization of glucose, xylose, and arabinose mixtures by engineered Saccharomyces cerevisiae strains. Appl. Environ. Microbiol. 75:907–914.

Yona AH, Manor YS, Herbst RH, Romano GH, Mitchell A, Kupiec M, Pilpel Y, Dahan O. 2012. Chromosomal duplication is a transient evolutionary solution to stress. Proc. Natl. Acad. Sci. U S A 109:21010–21015.

